# SPHERE: A novel approach to 3D and active sound localization

**DOI:** 10.1101/2020.03.19.998906

**Authors:** V. Gaveau, A. Coudert, R. Salemme, E. Koun, C. Desoche, E. Truy, A. Farne, F. Pavani

## Abstract

In everyday life, localizing a sound source in free-field entails more than the sole extraction of monaural and binaural auditory cues to define its location in the three-dimensions (azimuth, elevation and distance). In spatial hearing, we also take into account all the available visual information (e.g., cues to sound position, cues to the structure of the environment), and we resolve perceptual ambiguities through active listening behavior, exploring the auditory environment with head or/and body movements. Here we introduce a novel approach to sound localization in 3D named SPHERE (European patent n° WO2017203028A1), which exploits a commercially available Virtual Reality Head-mounted display system with real-time kinematic tracking to combine all of these elements (controlled positioning of a real sound source and recording of participants’ responses in 3D, controlled visual stimulations and active listening behavior). We prove that SPHERE allows accurate sampling of the 3D spatial hearing abilities of normal hearing adults, and it allowed detecting and quantifying the contribution of active listening. Specifically, comparing static vs. free head-motion during sound emission we found an improvement of sound localization accuracy and precisions. By combining visual virtual reality, real-time kinematic tracking and real-sound delivery we have achieved a novel approach to the study of spatial hearing, with the potentials to capture real-life behaviors in laboratory conditions. Furthermore, our new approach also paves the way for clinical and industrial applications that will leverage the full potentials of active listening and multisensory stimulation intrinsic to the SPHERE approach for the purpose rehabilitation and product assessment.

## INTRODUCTION

Spatial hearing is a fundamental ability for humans and other animals. Accurate localization of sounds allows for the construction of maps of the environment beyond the limits of the visual field, guides head and eye orienting behaviour, plays a crucial role in multisensory integration, supports auditory scene analysis and can improve discrimination of auditory signals from noise. In everyday environments, spatial hearing is three-dimensional, multisensory and active. We estimate azimuth, elevation and distance of sounds. We perceive the visual context in which they occur and often also the event that generated them (Kumpik et al. 2019). Most importantly, to resolve perceptual ambiguity in sound localization, we explore the auditory environment with head and body movements (Andéol & Simpson, 2016).

When spatial hearing abilities are investigated in laboratory or clinical settings, however, most of these naturalistic aspects of spatial hearing are considerably constrained. In the laboratory, participants are typically presented with sounds that originate from a limited set of positions, typically varying only along the azimuthal plane (i.e., all having fixed elevation and distance; Haber et al., 1993; Grantham et al., 2007). Few studies manipulated azimuth and elevation jointly (e.g. Oldfield & Parker 1984; Wightman & Kistler 1999; Ahrens et al., 2019), and even fewer have modified azimuth, elevation and distance across trials (e.g. Haber et al., 1993; Brungart et al., 1999). In addition, although sounds were delivered in 3D -- hence, supposedly perceived in 3D space -- the response was often limited to one dimension (e.g., in studies using a rotating dial to indicate angular sound position; Haber et al., 1993) or two dimensions (e.g., in studies using a remote display to indicate sound direction; Andéol et al., 2014). These approaches limit our understanding of perceived auditory space, and might overestimate sound localization performance by reducing stimulation and response uncertainty.

Multisensory contributions to sound localization are also often poorly controlled. Some studies allow full vision of the loudspeakers (e.g., van Hoesel & Tyler 2003), whereas others hide loudspeakers behind visible barriers (e.g., Nava et al., 2009; Pavani et al., 2001; Pavani et al., 2003). Yet, any visible barrier provides relevant cues for estimating sound elevation and distance. When visual cues are completely eliminated by placing subjects in total darkness (Pavani et al., 2008), the listening experience lacks entirely the multisensory context that characterizes everyday auditory environments. In addition, whenever participants are blindfolded (Ahrens et al., 2019), or are required to keep their eyes closed (Brungart et al., 1999), eye-position cannot be measured. But in natural conditions, eye-orienting responses permit encoding of sound position in retinocentric coordinates (Bulkin & Groh 2006; Pavani et al., 2008) and allow for associations between heard sounds and plausible visual sources. In addition, static and dynamic eye-position influence sound localization (e.g., Lewald & Ehrenstein 1996; Groh & Sparks 1992; Pavani et al., 2008).

Most importantly, to ensure reproducibility of sound characteristics across trials, conditions and experimental sessions, it is common practice to constrain head-movements in sound localization studies. Participants are required to place their head on a chin-rest (e.g., Brungart et al., 1999; Pavani et al., 2008) or keep their head still during each trial (Távora-Vieira et al., 2015; Litovsky et al., 2009; van Hoesel & Tyler 2003). Although the importance of head-movements for sound localization has been emphasized since the 1940s (Wallach, 1940), the contribution of spontaneous head-movement has been largely overlooked. Until recently, researchers have tracked head-movements more as a way to collect behavioural measures for sound localization accuracy (Slattery & Middlebrooks, 1994), rather than a way to study spontaneous head-movements during sound localization (for notable exceptions see Brimijoin et al., 2010; 2012).

In sum, currently dominant approaches to sound localization typically do not stimulate nor measure responses in 3D auditory space, cannot fully control for multisensory information and, crucially, do not allow for active listening through head-movements. Here, we report on a novel approach that circumvents all these limitations and allows deeper insights into spatial hearing by using spatially reproducible free-field sounds, full control over visual stimuli and unconstrained, yet real-time controlled, head-, eye- and hand-movements.

Starting from the pioneering work by Brungart and colleagues (1999), we have implemented a method to guide a real loudspeaker to precisely pre-determined head-centered coordinates in each trial. We combined real-time tracking of the loudspeaker’s position with continuously monitoring of the listener’s head-position. To this aim, a head-mounted display (HMD) worn by the participant allowed to determine speaker position in each trial. Furthermore, we took advantage of the HMD to measure eye-movements, and to achieve complete control over the available visual stimulation. Using virtual reality (VR) we also instructed participants as to desired postures (e.g., straight ahead orienting at the beginning of each trial). Finally, we tracked in real-time hand motion and obtained full description of hand kinematic responses to sound source position in 3D. We provide full-detail description of this new method -- named SPHERE -- and report on one experiment testing 3D sound localization in normal hearing participants. To examine the validity of our active listening approach we compared one condition in which participants were required to remain still during the sound delivery (static listening condition), with a condition in which they were free to move (active listening condition). We predicted improved sound localization performance in the active compared to static listening (Brimijoin et al., 2010; 2012). Our results provide validation of the SPHERE system and paradigm, additionally showing the contribution of head-movements to sound localization accuracy.

## METHODS

### Participants

20 participants (mean age = 46, SD = 18; 12 females; 18 right-handed) were recruited through advertisement (e-mail or flyer) to take part in the study. All reported normal or corrected-to-normal vision, and no history of hearing deficits. Participants were informed that they were about to participate in a sound localization study, that would require wearing a virtual reality HMD, and that their task was to localize as accurately as possible a sound delivered in the space around them using a hand-pointing response. If they agreed to participate, they were asked to sign the informed consent documents. The study was approved by the Comité Ethique d’Evaluation de l’Inserm (IRB00003888), and was conducted in accordance with the ethical standards of the 1964 Declaration of Helsinki.

#### Apparatus and stimuli

A schematic description of the SPHERE apparatus is shown in Figure 1. The left side of the figure illustrates equipment in the control room, whereas the right side illustrates equipment in the testing room.

**Figure 1.**
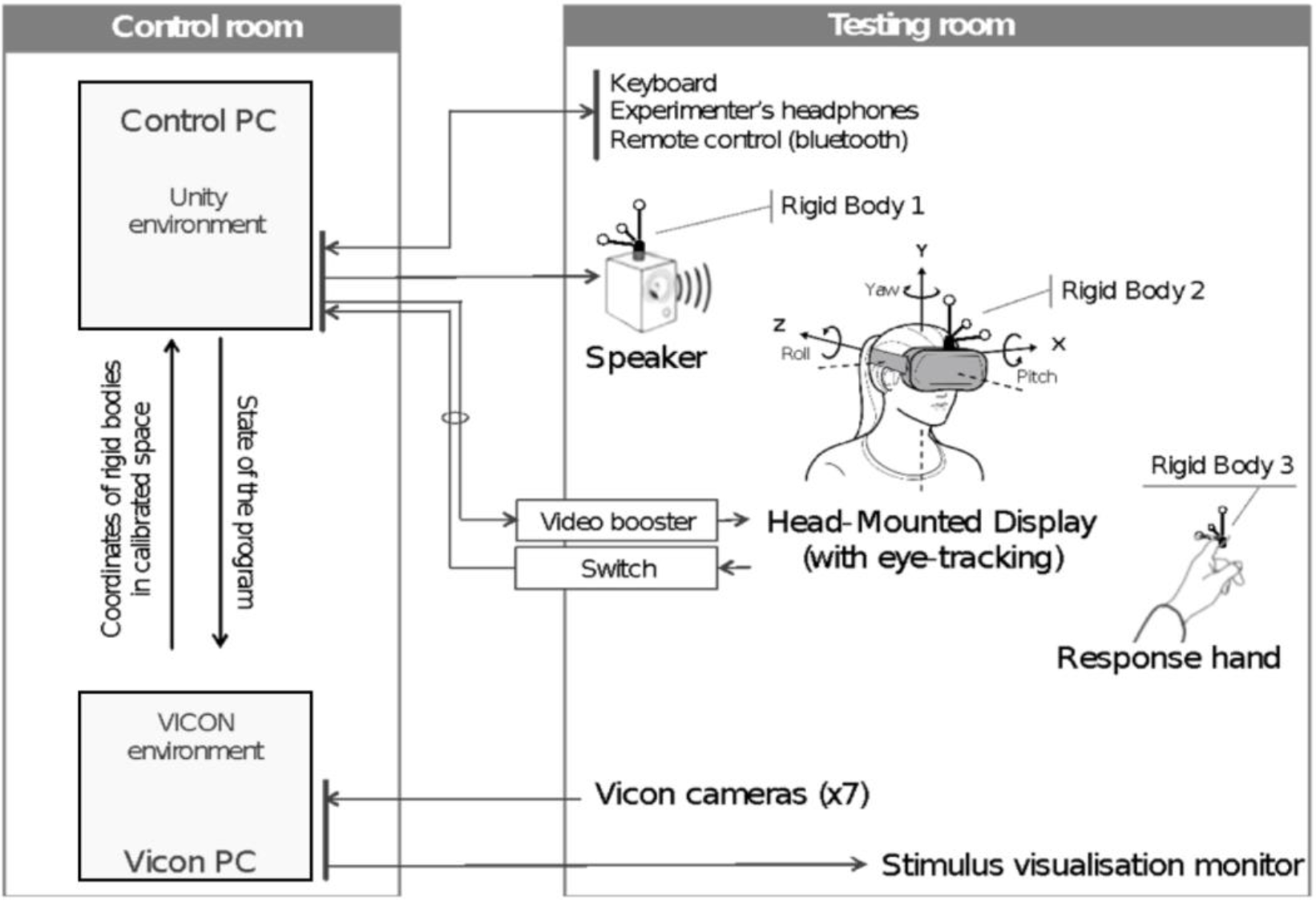
Schematics of the apparatus used for the study.

The control room hosted two desktop computers. The first computer, from now on ‘Control PC’, was an HP Z820 Workstation (Windows 7 Professional, Processor Intel(R) Xeon(R) CPU E5-2609 @ 2.40 GHz 2.40 GHz), equipped with a NVIDIA Quadro K5000 graphic card (DirectX 11.0). It controlled the entire sequence of events, stimulations, response collections and data saving through a custom-made script written in Unity (Version 5.5.1f1 32-bit, Unity Technologies, San Francisco, CA). The second computer, hereafter ‘Vicon PC’, was an HP Z230 Tower Workstation (Windows 7 Professional, Processor Intel(R) Core(TM) i7-4771 CPU @ 3.50 GHz 3.50 GHz). It controlled the Vicon motion capture system (Vicon Tracker 2.0.1 x64, Vicon Motion Systems LTD, Oxford, UK), and it ran a custom-made script written in Unity that served for stimulus visualization (see below).

The setting comprised Vicon cameras for motion capture, three rigid bodies for real-time object-tracking, the head-mounted display (HMD) incorporating eye-tracking system, one monitor for stimulus visualization, one loudspeaker, one keyboard and one remote control. Each of these pieces of equipment is described below, with details concerning the way it was interfaced with the Control and Vicon PCs.

##### Vicon motion capture

The Vicon motion capture system comprised 7 infra-red cameras (Bonita 10: Frame rate 250 fps, Resolution 1024×1024, Vicon^®^, Oxford, UK) mounted on the walls of the testing room. The elevation (195-205 cm) and semi-circular arrangement of the cameras allowed full kinematic tracking of a wide 3D space (height: 250 cm; width: 320 cm; depth: 170 cm). The space visible to the cameras was calibrated using the Vicon Active Wand tool (www.vicon.com/products/vicon-devices/calibration), which allows a multi-plane video calibration across the entire acquisition volume. Once calibrated, object-tracking spatial precision was <1 mm (down to 0,5 mm in a 4 × 4 meters volume). Then we placed the HMD on the floor in a straight-ahead position to recorded a straight-ahead reference direction (taking into account HMD rotations). The cameras were connected to a multiport box in the testing room, which in turn was USB connected to the Vicon PC in the control room.

##### Rigid bodies

The Vicon system captured the position of three distinct rigid bodies (each mounting 4 reflective 9 mm markers), with a sampling frequency of 100 Hz. The first rigid body (rigid body 1; radius 75 mm) was fixed on top of the loudspeaker and served for tracking its xyz coordinates in the calibrated space; the second rigid body (rigid body 2; radius 75 mm) was fixed on top of the HMD and served for tracking HMD and the head center positions; the third rigid body (rigid body 3; radius 75 mm) served for head-size calibration and for collecting hand-pointing responses.

##### Head-mounted display

The HMD was an Oculus Rift Development Kit 2 system (DK2, Oculus VR^®^, Menlo Park, USA. Screen OLED, Resolution: 1920 x 1080 (960 x 1080 per eye), maximal refresh of 75Hz, dimensions L x W x H: 1.3 x 14.7 x 7.1 inches, and a field of view equal to 106°) running with Oculus Runtime (Version 0.6; Facebook Technologies Ireland, Dublin, Ireland). The Oculus Rift DK2 incorporated an eye-tracking system (SensoriMotoric Instruments, Berlin, Germany; www.smivision.com; 60Hz frequency and 0.5 degrees spatial precision). In our setup, the HMD served two purposes: (1) it conveyed visual instructions to the participant; (2) it allowed continuous monitoring of the participant’s eye movements.

##### Loudspeaker

.A loudspeaker (JBL GO Portable, 68.3 x 82.7 x 30.8 mm, Output Power 3.0W, Frequency response 180 Hz – 20 kHz, Signal-to-noise ratio > 80 dB) was used to deliver all target-sound stimuli. Target stimuli were amplitude modulated broad-band bursts lasting 3 seconds (see Supplementary Figure S1 for spectrograms).

A keyboard, a remote control (Targus^®^, Laser Presentation Remote) and a monitor (DELL 19’’ 5:4, resolution 1280 x 1024), completed the equipment in the testing room. All devices were connected to the control PC, except the stimulus visualization monitor that streamed a copy of the screen of the VICON PC to be seen inside the testing room. The function of these four pieces of equipment is described in details in the procedure section below.

#### Procedure

Before starting the experiment, participants were introduced to the task and to the virtual reality equipment using a picturized information sheet and a custom-made video that showed one experimenter wearing the HMD and performing several sound localization trials. It was made explicit to all participants that their task was to listen carefully to the sound and indicate its location in space using the pointer held in their hand (i.e., rigid body) after the sound terminated.

Participants were told that sounds could be delivered everywhere in the 3D space around them (e.g., above and below ear level, in front and back space, at multiple azimuthal positions). They were also informed that all sounds would be delivered within their arm reaching distance. They were also told they would perform the sound localization task under two conditions. One (static listening) in which they would have to keep their head still in the initial position throughout sound presentation and another condition (active listening) in which they were prompted to actively search for the sound during its presentation, by freely moving their head. In both conditions, they were free to move their head and body as soon as the sound terminated (e.g., rotation on the chair to reach for sounds in back space). After receiving the instructions, participants were conducted in the testing room, seated on a rotating office chair (the center of the head being aligned with the center of the chair rotation axis), and prepared for the task with the HMD and the hand-held rigid body.

The experiment began with eye- and head-center calibrations: (i) Eye-calibration was performed using a 5-points calibration grid (smart recorder of SMI Eye tracking software) and it permitted control of the 3D cyclopean eye position and direction; (ii) Head-center calibration was performed by collecting the 3D position of the two ears (using rigid body 3), averaging these positions to obtain the 3D head center position. The head-center position served as origin for the polar coordinate system that included speaker, hand, head and cyclopean gaze positions. Both eye and head calibrations were carried out each time the HMD was displaced (e.g., when participant took breaks during the experimental session).

The loudspeaker position in 3D space was calculated with reference to the center of the head. In this way, despite participants sat without any chin-rest, we could carefully control the position of each sound source with respect to the ears. Twelve pre-predetermined positions were used throughout the experiment, resulting from the combination of 4 different azimuths (−30°, 30°, −150° or 150°), 3 different depths (35 cm, 55 cm or 75 cm) and a single elevation (0°, i.e., ear-level). Figure 2 shows a bird-eye view of the 12 positions around the participant’s head.

**Figure 2.**
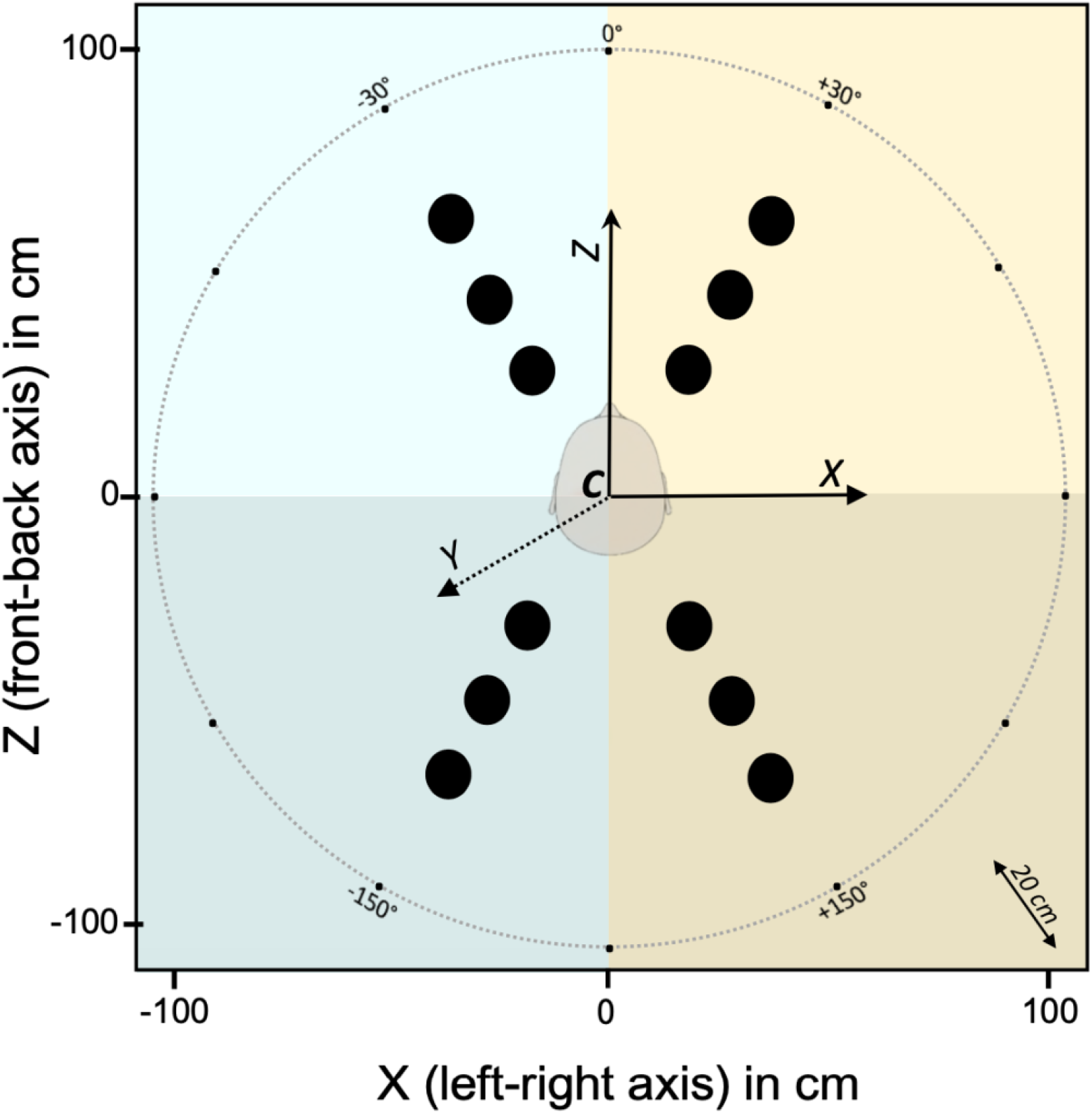
Bird-eye view of all pre-determined loudspeaker positions in head-centered coordinates. The origin of the polar reference-frame used for sound positioning was the participant head-center, measured in real-time (marked by C in the image). The 3D axes of this reference frame were the azimuth (X), the elevation (Y) and the depth (Z). Twelve pre-determined loudspeaker locations (white circles) were arranged around the participant’s head: at 4 different azimuths (X-values: ±30° and ±150°), 3 different depths (Z-values: 35, 55 or 75 cm), all at ear-level (Y-values: 0°).

In each trial, two sets of instructions generated in real-time by the computer informed the experimenter as to where to position the loudspeaker in the 3D space surrounding the participant (i.e., one of the 12 predetermined positions detailed above). The stimulus visualization monitor (see Figure 1) displayed in real-time the actual position of the loudspeaker and its desired position for the upcoming trial. This allowed the experimenter to rapidly reach for each computer-determined position, keeping the speaker at ear-level. Information about the precise elevation positioning of the speaker was given to the experimenter via one in-ear headphone (i.e., non-audible by the participant) and consisted in an echo radar sound, which increased in frequency and intensity as the speaker approached the target position. The system software considered the loudspeaker to be correctly positioned when it entered a sphere of 5 cm diameter centered on the pre-determined sound position and delivered the sound only when three criteria were concurrently met: (1) the loudspeaker was in the 3D position pre-determined for the trial; (2) the participant’s head was facing straight ahead; (3) the participant’s eyes were directed straight ahead. Participants actively complied with criterion 2 and 3 by aiming their head and eyes to align two crosshairs displayed in the HMD. Figure 3 shows the instructions visualized in the HMD (bottom part of each panel) and a cartoon of the participant’s corresponding head- and eye-posture (upper part of each panel), in the phases that preceded sound delivery. As shown in Figure 3A, at the beginning of each trial two white crosshairs were presented to the participant: a bold cross with a ball inside, indicating the desired position of the head and eyes, respectively. A thinner cross provided participant with visual feedback of their actual head-position. Participants were instructed to move their head to align the two crosshairs. When the alignment was achieved, the bold cross turned blue (Figure 3B). Likewise, participants were instructed to gaze the ball inside the bold cross. When fixation was achieved, the ball turned blue (Figure 3C). Once all the three criteria were met, all visual stimuli disappeared (i.e., the HMD display turned black) and sound emission started (Figure 3D). Participants were instructed to respond only after sound emission ended, by bringing the hand-held rigid body where they perceived the sound to originate and hold it there for two seconds. The experimenter validated this position by pressing a button on the remote control, which also terminated the trial. After trial completion, no feedback on performance was ever provided. Each trial lasted approximately 10-15 seconds, with the speaker positioning phase lasting about 3-5 seconds, depending on the pre-determined position. Recall that multiple aspects contributed to trial duration: events before sound delivery (the participant actively moved head and eyes to the desired initial posture, the experimenter manually brought the loudspeaker to the pre-determined position), the sound delivery itself, and the participant’s full-body motor response to the target after sound delivery.

**Figure 3.**
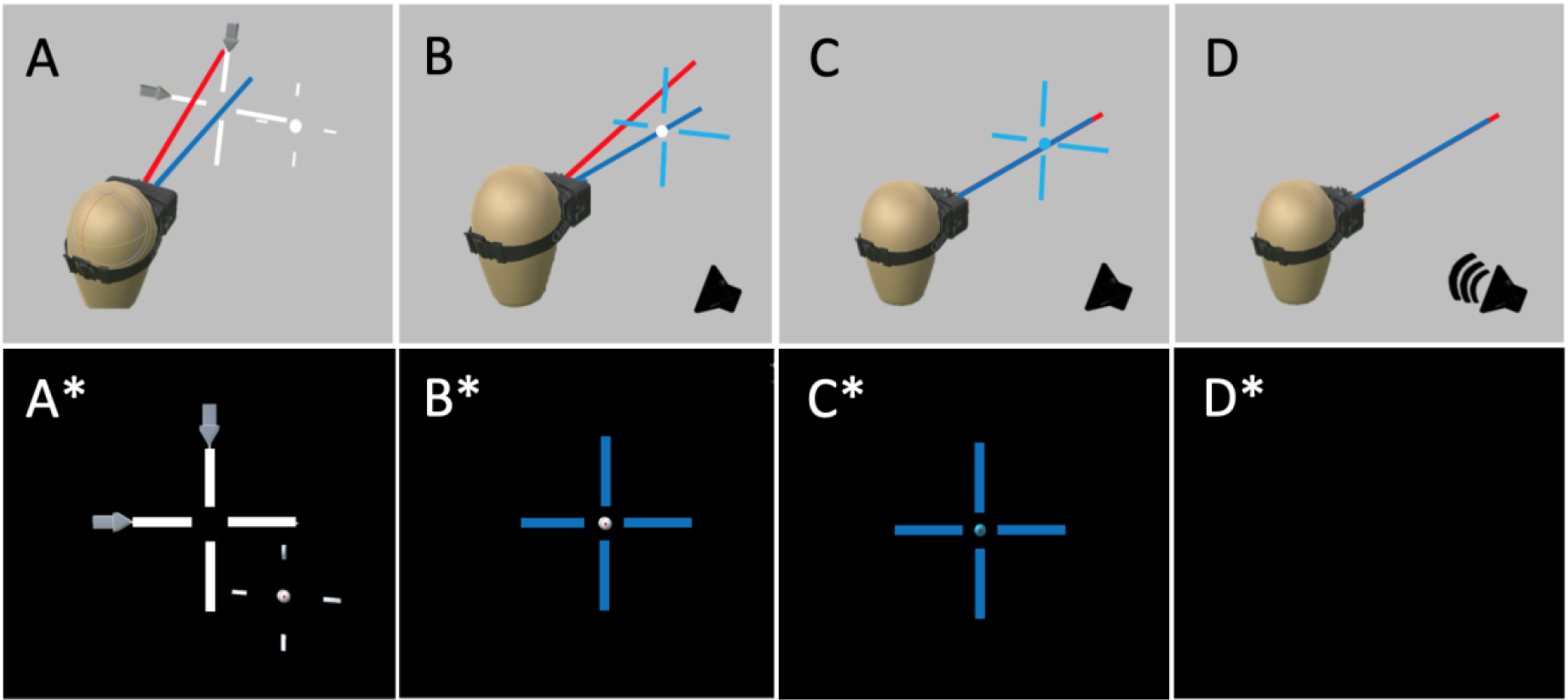
Pre-stimulation alignment of head and eyes. (A) At the beginning of each trial, the participant wearing the HMD was free to move is head (symbolized head direction: blue line) and eyes (symbolized cyclopean eye direction: red line). (A*) In the HMD display, the participant saw a bold white cross (indicating the actual position of the head), and a thin white cross with a central ball (indicating the desired position of the head and eyes, respectively). Two grey arrows flanked the bold white cross, showing the participant in which direction to move his head to achieve the desired initial position (in this example, the participant had to move his head to the right and down to ensure alignment). (B*) When the desired head-position was achieved the bold cross turned blue. (C*) When the desired eye-gaze position was reached, the central ball turned blue. (D) When all criteria were met (i.e., head-position straight ahead, eye-position straight ahead, and speaker within a sphere around the pre-determined position) all visual stimulations were removed, (D*) the scene became entirely dark, and the sound was delivered.

The experimental session was organized in 4 successive blocks, with a pause between each block in which the HMD was removed. Listening conditions (static or active) changed between blocks of trials. Half of the participants followed an Active-Static-Static-Active sequence, whereas the other half followed a Static-Active-Active-Static sequence. Each block comprised 48 randomized trials (i.e., 4 trials for each of the 12 pre-determined positions), resulting in a total of 192 trials (i.e., 8 trials for each predetermined position in each listening condition). The entire experimental protocol lasted approximately 40 minutes.

#### Data processing

The position of all tracked elements recorded in Vicon reference frame (loudspeaker, head center and direction, hand) was re-computed in head-center reference frame, kinematic analysis of head and hand were analyzed and inspected for each trial by a custom-made software running on MATLAB R2013a. 3D position signal was filtered (50 Hz cut-off frequency, finite impulse response filter FIR) and velocities were computed from the filtered position signal using a two-point central difference derivative algorithm (Bahill & McDonald, 1983). In order to determine the sequence of head and hand movements, the beginning and the end of all movements were automatically detected using a velocity threshold procedure (80 mm/s). The results of this automatic procedure were then inspected off-line and corrected manually, if necessary. This procedure served to obtain the spatio-temporal profile of head and hand behavior, and to extract relevant parameters for subsequent analyses (number of head movements during sound emission, onset of the first head movement, onset of the head and hand response). It also served to reject all trials in which participants did not comply with the instructions (i.e., they made anticipatory hand movement during sound delivery, or head movement erroneously produced in the static listening condition).

An example of this procedure for a correctly executed trial in the active listening condition is shown in Figure 4A and 4B, for head and hand, respectively. In this example, one single head movement is visible during sound delivery (Figure 4A), and the head and hand response movements are clearly identified in the response phase (Figures 4A and 4B).

**Figure 4.**
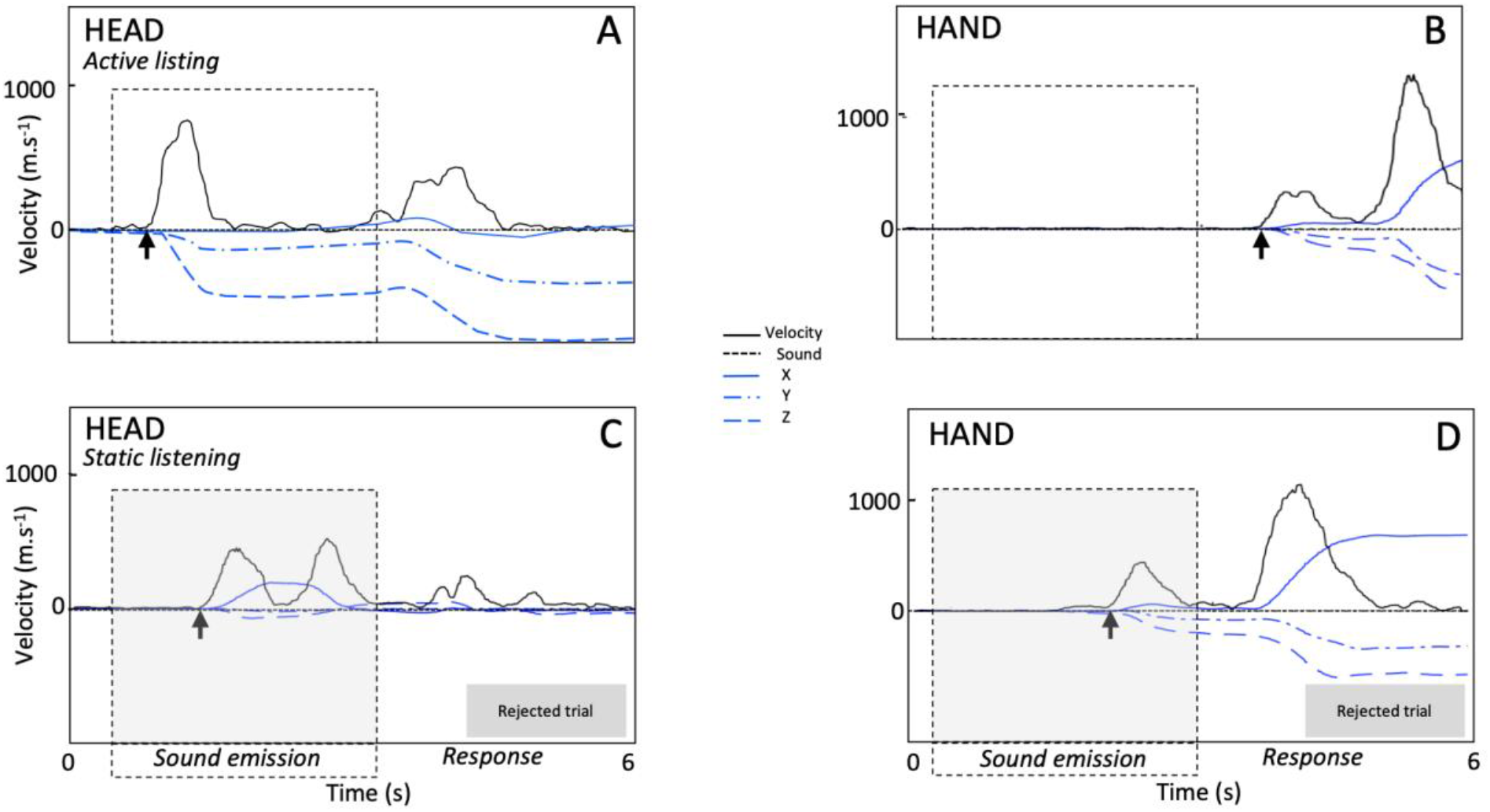
Description of spatio-temporal behavior of the head (A, C) and the hand (B, D), as shown by our custom-made software, during representative trials. The dotted black line shows the time of sound emission, the bold black line and the blue lines respectively shows the velocity profile and the x, y, z positions of the tracked element (head or hand). The arrow marks the beginning of the head or hand movement. Examples illustrate correctly executed trials (A and B), and rejected trials in which instructions were violated (C: head movement initiated during sound emission in a static listening trial; D, hand movement initiated during sound emission).

An example of this procedure for two rejected trials is shown in Figures 4C and 4D. In Figure 4C, two head movements are visible during sound delivery in a trial which required static listening. In Figure 4D, the hand moves during sound emission.

All statistical analyses and data visualizations were performed using R and R-studio environment. Unless otherwise indicated, means ± standard errors are reported in the text. We planned ANOVAs or t-test, and the Greenhouse-Geisser sphericity correction has been applied to analyses of variance, when appropriate. For all post-hoc comparisons, the FDR (False Discovery Rate) method implemented in R (Benjamini & Hochberg, 1995) has been adopted to correct for multiple comparisons when needed.

## RESULTS

### Positioning of speakers at pre-determined locations

The apparatus we developed poses minimal constraints on the participants’ body movements: movements were unrestrained in all three dimensions and, after sound delivery, participants were free to rotate on the chair to explore sounds in back space. Yet, at the beginning of each trial, participants were instructed to align their head and eye with respect to straight ahead position. On-line kinematic tracking allowed to deliver sounds only when the required eye and head posture criterions were matched.

Moreover, the participant’s compliance with the instructions was further examined through off-line kinematic analyses using a custom-made software. Trials in which instructions were not followed were excluded from further analyses (static listening: 6.1%, SD = 8; active listening: 6.6%, SD = 7). The main reason for trial rejection was anticipatory hand responses.

The absence of physical constraints on the participants’ posture poses potentially a problem of reproducibility of sound source position across experimental trials and participants. This because the head returns always to different initial positions after the response (hand-pointing to sounds delivered all around the participant). In the next paragraphs, we show how head-centered positioning of the loudspeaker effectively solved this issue.

We started by testing if the pre-determined loudspeaker locations remained constant across trials and participants. Figure 5 shows initial head position (in black) and 12 pre-determined locations (in gray) for all participants across 192 trials in VICON reference frame (Fig. 5A and C, in bird-eye and lateral view respectively). A substantial variability is observed, which however reduces dramatically when all positions are referenced to head-center, i.e., converted to a head-centered reference frame (Fig. 5B and D). Figure 5E summarizes the effect of head-referencing by comparing mean changes in standard deviation (with standard errors) across participants in VICON vs. head-centered coordinates (x: 1.08 ± 0.24 cm vs. 0.10 ± 0.02 cm; y: 0.63 ± 0.14 cm vs. 0.10 cm; depth: 1.89 ± 0.42 cm vs. 0.15 ± 0.03 cm)

**Figure 5.**
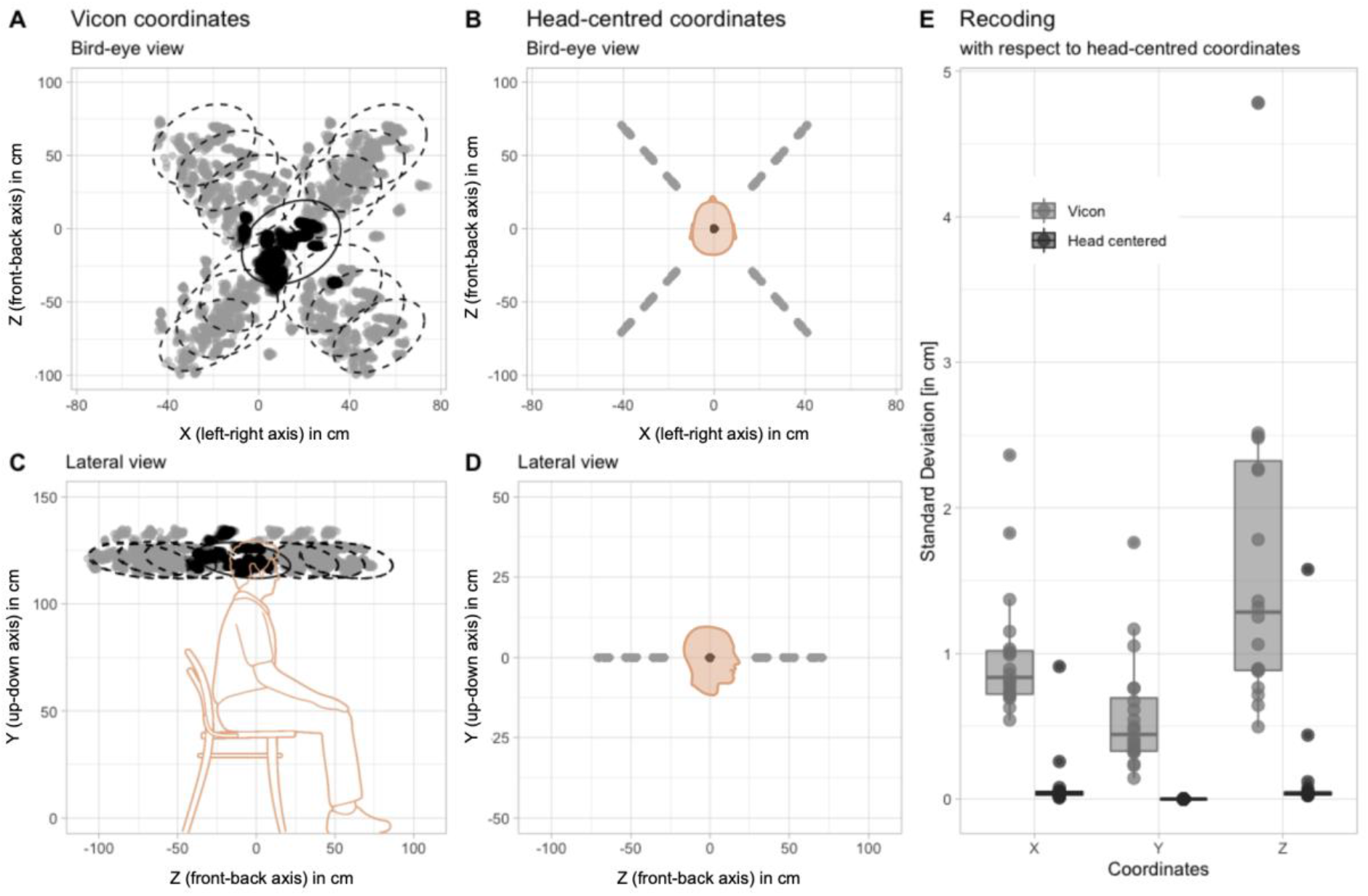
Normalization to head-centered coordinates. Bird-eye and lateral view of initial head position (in black) and 12 pre-determined locations (in gray) for all participants (192 trials each), in VICON reference frame (A-C) and head-centered coordinates (B-D). Variability of pre-determined locations averaged across 12 positions for each participant in the two reference frames, as a function of coordinates x, y, z (E).

Next, we tested if the variability of loudspeaker actual location around the predetermined position was within the established tolerance (i.e., a sphere with a 5 cm diameter around the pre-determined position). Figure 6 shows all 192 stimulations for all participants, when the 12 pre-determined locations are re-aligned to a single coordinate, centered on the origin of the axes. Dashed ellipses in the 2D panels of Figure 6 represent 95% confidence interval of the distribution, and indicate that for all these trials stimulation remained within 2.5 cm from the pre-determined location (mean differences with SE between pre-determined and actual location: x = 0.87 ± 0.01 cm, y = 1.15 ± 0.02 cm, z = 0.93 ± 0.01 cm; error in 3d = 1.98 ± 0.01 cm). (mean differences between pre-determined and actual location: x = 0.87 ± 0.6 cm, y = 1.15 ± 0.8 cm, z = 0.93 ± 0.7 cm).

**Figure 6.**
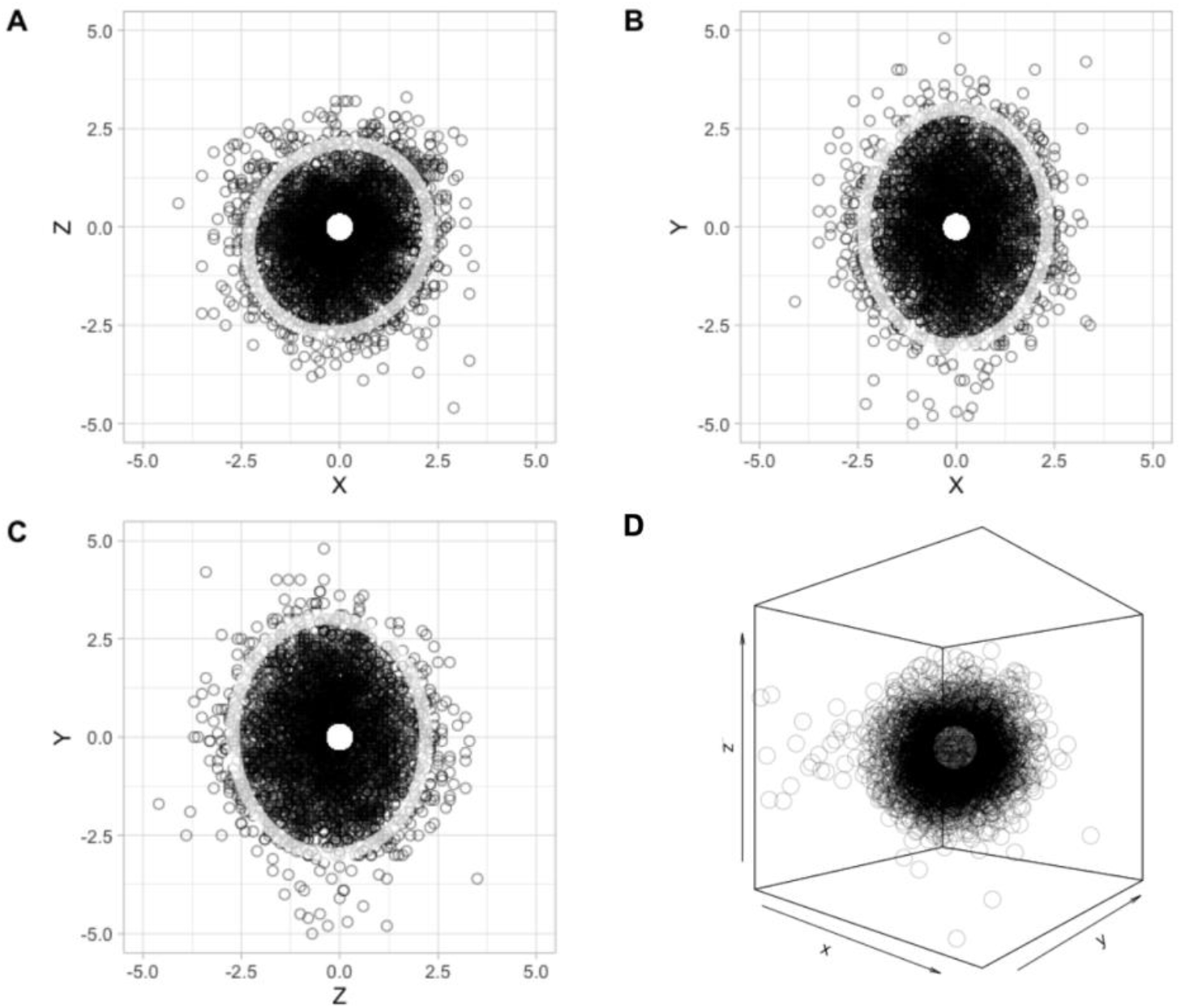
Actual speaker location with respect to pre-determined location, in centimeters. Stimulations delivered to all participants, when the 12 pre-determined locations are re-aligned to a single coordinate, centered on the origin of the axes. (A) top view; (B) front view; (C) lateral view; (D) 3D rendering. Ellipses in the 2D panels represent 95% confidence interval of the distribution.

### Hand pointing to sounds in static listening

Sound localization performance during static listening is shown in Figure 7, for azimuth, elevation and distance separately. Dispersion plots show hand-pointing responses for all trials and participants (with 95% confidence intervals shown by ellipses), color-coded as a function of sound distance (far, middle and near). Bar plots summarize means and 95% confidence intervals for absolute and variable errors in each of the three dimensions (for performance details, see Table 1).

**Figure 7.**
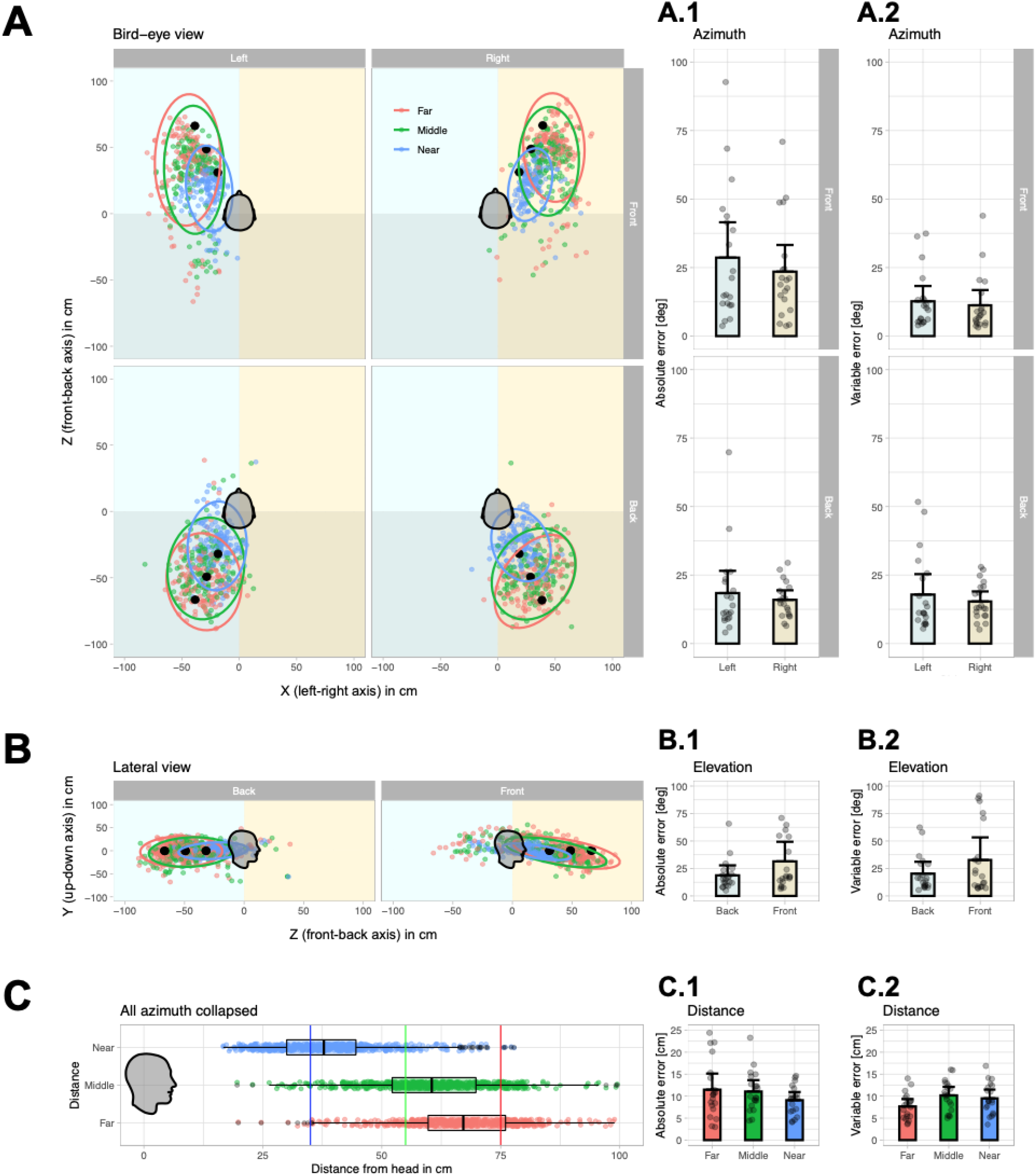
Sound localization performance during static listening. (A) Bird-eye view of all target positions (black dots) and hand-pointing responses (colored dots) in all trials and participants. Ellipses indicate 95% confidence interval of responses within each quadrant and distance. Responses and ellipses are color-coded as a function of target distance: red for far, green for middle, and blue for near targets. Absolute (A1) and variable (A2) error in azimuth for each participant (dots), as a function of side (leftright) and antero-posterior position (front-back) of target sounds. Error bars indicate the 95% confidence intervals, adjusted for within-subjects’ designs using method from Morey (2008). (B) Lateral view of all target positions and responses of all participants. Absolute (B1) and variable (B2) error in elevation for each participant (dots) as a function of antero-posterior position of target sounds. Responses for each participant are averaged across side (left or right) and distance. (C) Lateral view of responses (with boxplots) as a function of sound distance. Absolute (C1) and variable (C2) errors in depth for each participant (dots).

**TABLE 1.** Mean absolute, signed and variable errors for the three dimensions separately (azimuth, elevation and distance), and for all dimensions (3D-error). The mean errors and standard deviation were calculated separately for each participant and 12 sound positions, then combined to generate the values in the table. Mean errors were segregated in respect to listening postures (Static and Active posture), front and backspace, and left and right sound sources. The errors in azimuth and elevation are reported in degrees, and the distance errors are in centimeters, the 3D-error is a computation of all dimension errors (see paragraph ‘3D-error’). (standard errors were put in parenthesis).

|  | Azimuth (in degrees) |  |  | Elevation (in degrees) |  |  | Distance (in centimetre) |  |  | 3d error (in cm) |
| --- | --- | --- | --- | --- | --- | --- | --- | --- | --- | --- |
|  | Absolute error | Signed error | Variable error | Absolute error | Signed error | Variable error | Absolute error | Signed error | Variable error | D |
| **Static posture** |  |  |  |  |  |  |  |  |  |  |
| Front | Mean (SE) | Mean (SE) | Mean (SE) | Mean (SE) | Mean (SE) | Mean (SE) | Mean (SE) | Mean (SE) | Mean (SE) | Mean (SE) |
| Left | 28.6 (5.3) | -28.2 (5.4) | 12.7 (2.3) | 34.2 (8.3) | -2.9 (7.9) | 30.4 (8.7) | 10.0 (0.7) | -5.4 (1.6) | 8.7 (0.5) | 34.7 (4.3) |
| Right | 23.5 (4.0) | 23.2 (4.1) | 11.2 (2.3) | 28.4 (6.1) | -5.1 (7.2) | 23.5 (6.1) | 9.6 (0.7) | 2.0 (1.7) | 9.6 (0.6) | 32.6 (3.8) |
| Mean | 26.1 (4.0) | -2.4 (2.6) | 29.4 (4.4) | 31.4 (6.0) | -4.2 (6.9) | 32.7 (6.9) | 9.8 (0.6) | -1.7 (1.5) | 10.2 (0.6) | 34.5 (3.7) |
| Back |  |  |  |  |  |  |  |  |  |  |
| Left | 18.4 (3.4) | 5.8 (4.3) | 17.9 (3.1) | 20.7 (5.0) | 4.7 (5.7) | 20.4 (4.5) | 11.6 (1.4) | -3.8 (2.4) | 10.4 (0.7) | 27.1 (2.2) |
| Right | 16.0 (1.4) | -6.0 (2.7) | 15.4 (1.5) | 17.3 (2.0) | -1.6 (4.0) | 16.2 (2.2) | 10.9 (0.8) | 1.4 (1.8) | 11.5 (0.7) | 28.5 (1.8) |
| Mean | 17.1 (2.2) | 0.2 (2.2) | 20.5 (2.7) | 18.7 (3.1) | 1.6 (4.3) | 20.3 (3.6) | 11.3 (1.0) | -1.4 (2.0) | 11.5 (0.7) | 27.9 (1.7) |
| Overall | 21.6 (2.3) | -1.1 (2.2) | 27.6 (2.9) | 25.1 (3.4) | -1.3 (3.4) | 35.5 (5.0) | 10.5 (0.7) | -1.5 (1.6) | 11.3 (0.6) | 32.2 (2.3) |
| **Active posture** |  |  |  |  |  |  |  |  |  |  |
| Front |  |  |  |  |  |  |  |  |  |  |
| Left | 18.5 (5.0) | -17.9 (5.1) | 8.1 (1.3) | 21.8 (6.3) | 1.1 (7.7) | 12.5 (1.9) | 10.0 (0.8) | -2.9 (1.8) | 9.2 (0.4) | 27.1 (3.5) |
| Right | 13.3 (3.0) | 11.8 (3.1) | 9.5 (2.4) | 20.5 (4.6) | 2.2 (5.5) | 14.7 (4.0) | 10.0 (0.9) | 3.7 (1.7) | 10.0 (0.6) | 27.4 (3.6) |
| Mean | 16.0 (3.8) | -3.2 (1.8) | 18.3 (4.2) | 21.2 (5.2) | 1.6 (6.4) | 15.4 (3.3) | 10.0 (0.6) | 0.4 (1.7) | 10.4 (0.5) | 27.6 (3.5) |
| Back |  |  |  |  |  |  |  |  |  |  |
| Left | 17.8 (3.1) | 2.9 (4.3) | 16.9 (3.5) | 21.7 (4.6) | -2.7 (4.5) | 21.6 (5.8) | 11.6 (1.4) | -5.0 (2.4) | 10.6 (0.8) | 28.0 (2.3) |
| Right | 15.1 (2.0) | -6.4 (3.1) | 12.3 (1.3) | 17.0 (2.3) | -4.5 (3.8) | 16.0 (2.7) | 10.0 (0.8) | 0.3 (1.7) | 10.3 (0.7) | 27.9 (2.1) |
| Mean | 16.7 (2.4) | -1.6 (2.3) | 20.0 (3.1) | 19.7 (3.1) | -3.5 (4.1) | 21.3 (4.3) | 10.9 (1.0) | -2.5 (1.9) | 11.3 (0.7) | 28.4 (2.0) |
| Overall | 16.2 (2.5) | -2.3 (1.9) | 20.9 (3.0) | 20.2 (3.2) | -0.8 (2.8) | 26.3 (4.0) | 10.4 (0.7) | -1.0 (1.7) | 11.3 (0.5) | 28.5 (2.4) |

The bird-eye view in Figure 7A shows that the vast majority of trials fell within the correct quadrant and were distributed radially along the stimulation axis. Responses to front sounds were more eccentric than actual sound position, whereas this pattern was not evident for back sounds. To confirm this observation statistically, we entered the absolute (Figure 7A1), variable (Figure 7A2) and signed error (not shown) in the azimuth dimension into separate analysis of variance (ANOVA) with side (left and right) and antero-posterior sector (front and back) as within-participants factors. The analysis on signed error revealed a two-way interaction (F(1,19) = 54.60, p < .001, 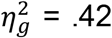) that confirmed larger lateral biases in pointing responses in front compared to backspace, for right (front: 23.2 ± 4.7 degrees; back: −6.0 ± 3.1 degrees; p < 0.001) and left target sounds (front: −28.2 ± 6.3 degrees; back: 5.8 ± 5.0 degrees; p < 0.001). The difference between front than back space also approached significance when absolute errors were considered (F(1,19) = 3.58, p = .074, 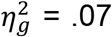). No other main effect or interaction reached or approached significance.

The lateral view in Figure 7B shows that minimal front-back confusions occurred (back to front = 2.6% ± 1.9; front to back = 10.7 ± 3.5) and that vertical dispersion was rather limited (variable error front: 32.7 ± 9.8 degrees; back: 20.3 ± 5.2degrees). To study performance in elevation as a function of stimulation quadrant we entered absolute, variable and signed error into separate ANOVAs similar to one described above. Although absolute (Figure 7B1) and variable (Figure 7B2) errors in elevation were numerically larger for targets in front compared to backspace only marginally significant differences emerged (absolute error, main effect for antero-posterior sector: F(1, 19) = 3.35, p = .083, 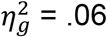).

Finally, Figure 7C shows that participants were able to distinguish the three target distances, yet with an average absolute error of 10.5 ± 1.3 cm. ANOVAs with target distance (near, middle and far) as within-participants factor, showed that target distance did not affect the absolute error (Figure 7C1), but significantly impacted on signed (F(1.15, 21.94) = 30.23, p < .001, 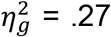) and variable errors in Figure 7C2 (F(1.54, 29.22) = 5.72, p = .013, 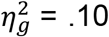). Participants underestimated far targets (−8.4 ± 2.5 cm) compared to middle (3.3 ± 2.5 cm; p<0.0001) and near targets (0.2 ± 1.7 cm; p<0.0001). In addition, dispersion of responses in depth was smaller for far targets (7.7 ± 0.8 cm) compared to middle (10.2 ± 0.9 cm; p<0.0001) and near targets (9.4 ± 1.0 cm; p<0.0001).

### Hand pointing to sounds in active listening

Having characterized sound localization responses in azimuth, elevation and depth in the static listening condition, we turn to investigate whether active listening (i.e., free head-movements during sound presentation) changes spatial hearing. Dispersion plots on the left side of Figure 8 show hand-pointing responses (averaged across trials delivered from each loudspeaker position, separately for each participant), color-coded as a function of listening condition (black: static; red: active). Line plots on the right side of Figure 8 depict changes in absolute localization errors between listening conditions, separately for each participant (the bold horizontal line indicates the mean).

**Figure 8.**
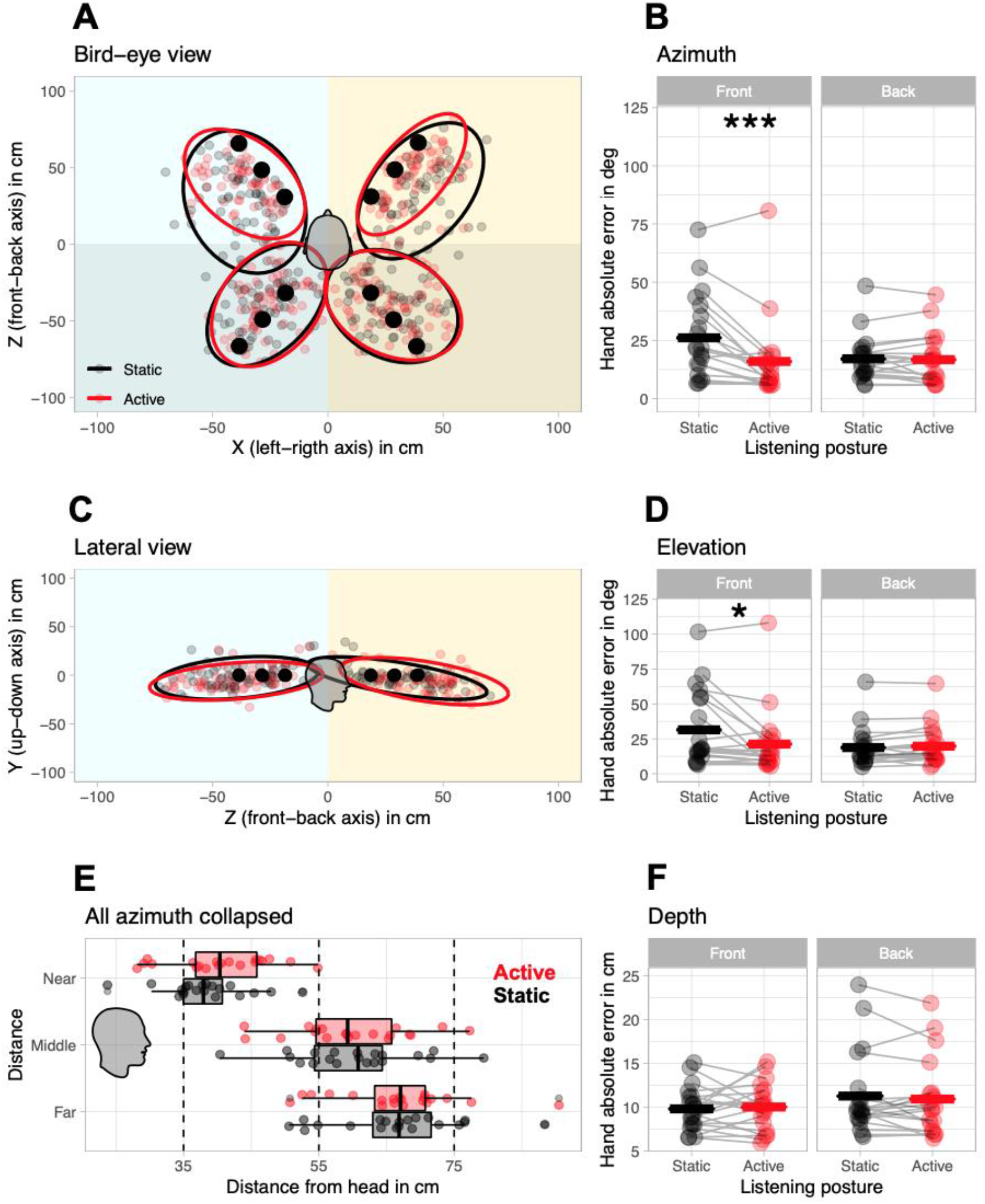
Effects of static and active listening on sounds localization. (A) Bird-eye view of all target positions (black dots) and hand pointing responses (smaller gray and red circles) for each participant, averaged across trials in a quadrant (i.e. front-left, front-right, back-left, back-right) irrespective of sound distance. Ellipses indicate 95% confidence interval of the responses across participants within each quadrant. Ellipses are color-coded as a function of listening condition (black: static listening; red: active listening). (B) Hand absolute error in azimuth for each participant, as a function of listening condition and antero-posterior position of target sounds. Bold horizontal lines indicate the mean for all participants. (C) Lateral view of all target positions and responses. Responses for each participant are averaged across side (left or right) and distance (near, middle or far). (D) Hand absolute error in elevation for each participant. (E) Lateral view of responses in depth (black boxplot: static listening; red boxplot: active listening). (F) Hand absolute error in depth for each participant. Asterisks indicate significant differences (* p<0.05; ** p<0.01; *** p<0.001).

The bird-eye view in Figure 8A shows that 95% confidence ellipses reduces in active compared to static listening, in front space selectively. To study azimuth errors along the antero-posterior axis as a function of listening condition, we entered absolute and variable errors in separate ANOVAs with antero-posterior sector (front and back) and listening condition (static and active) as within-participants’ factors. The analysis on absolute errors revealed the expected 2-way interaction (F(1, 19) = 12.93, p = .002, 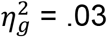), Azimuth absolute error in front space reduced for active (15.9 ± 3.8 degrees) compared to static listening (26.1 ± 4.0 degrees, p = 0.0005), whereas no such change occurred in back space (static: 17.2 ± 2.2 degrees; active: 16.5 ± 2.3 degrees; p = 0.4; Figure 8B). Instead, azimuth variable error reduced in active (11.7.9 ± 1.4) compared to static listening (14.3 ± 1.3), irrespective of whether stimuli were in front or back space (main effect of antero-posterior sector, F(1, 19) = 4.55, p = .046, 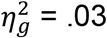).

A convergent result emerged for elevation (Figure 8C). When absolute and variable elevation errors were entered into an ANOVA similar to the one described above, the 2-way interaction between antero-posterior sector and listening condition reached significance for absolute errors (F(1, 19) = 6.54, p = .019, 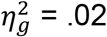). Absolute errors in front space reduced for active (21.2 ± 5.1 degrees) compared to static listening (31.4 ± 6.0 degrees, p = 0.02), whereas no such change occurred in backspace (static: 19.0 ± 3.1 degrees; active: 19.3 ± 3.0 degrees; p = 0.7; Figure 8D). Again, variable error reduced in active (16.2 ± 2.6) compared to static listening (22.6 ± 3.2), irrespective of whether stimuli were in front or back space (main effect of antero-posterior sector, F(1, 19) = 4.59, p = .045, 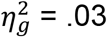).

By contrast, active listening did not affect depth estimation (Figure 8E). Absolute errors in depth were entered into an ANOVA with distance (near, middle, far), antero-posterior sector (front and back) and listening condition (static and active) as within-participants factors. No significant main effect or interaction involving listening posture emerged (all F < 2.06). Likewise, no main effect or interaction involving listening condition emerged for variable errors in depth.

### Head movements during active listening

Kinematic tracking of the HMD allowed detailed investigation of head movements during sound in the active listening condition. Despite active listening allowed free head-movements during sounds, participants moved their head in 66.6% of trials on average (SD = 31.4%; median = 75.7, IQR = [58.3 90.9]). In fact, the distribution of percent head-movements (Supplementary Figure S2A) revealed 2 outliers: one participant that never moved his head and another that moved only on 6 out of 96 trials (6.3%). These two outliers were removed from all subsequent analyses on head-movements.

The mean number of head movements during sound was 1.22 ± 0.04, with an average onset at 1077 ± 73 ms (head-movements beyond 3000 ms, i.e., after sound emission, were removed from this analysis; Figure S2B). Head-movements occurred both for targets in front and back space (75.7% and 72.9% of trials, respectively). For targets in front space they were mostly directed to the target. On average, for sounds at +30 degrees the first head movement was directed to 23.6 ± 1.8 degrees, whereas for sounds at −30 degrees it was directed to −27.0 ± 2.5 degrees (Figure S2C). For targets in back space head movements were distributed within the entire stimulated hemispace (Figure S2D). They were either directed to the front quadrant on the same side as the target (e.g., left front quadrant for targets at −150 degrees) or were aimed directly at the back target (in this case involving a trunk movement). On average, for sounds at +150 degrees the first head movement was directed to 107.5 ± 6.3 degrees, whereas for sounds located at −150 degrees it was directed to −118.0± 6.6.

### 3D-error

As a final step, we quantified overall sound localization performance in 3D (i.e., across azimuth, elevation and depth, and irrespective of sound position) and studied how this cumulative index was affected by listening condition. To this aim, we adapted the error system introduced by Rakerd & Hartmann (1986; see also Grantham et al., 2008), which combines into a single measure the absolute constant error (referred to as <u>C</u>) and the random error (referred to as <u>*s*</u>) as follows: 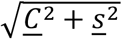. To obtain overall error in 3D we calculated in each trial *i*. the norm of the vector *<u>C</u>i*. (as shown in Figure 9A). This is the distance in 3D space between the participant’s response (i.e., the coordinates of rigid body held in the participant’s hand, *x_h_, y_h_, z_h_*) and the speaker location at the moment sound was delivered (i.e., the coordinates of the rigid body mounted on the speaker, *x_s_, y_s_, z_s_*). All *<u>C</u>i* extracted for each participant were then averaged irrespective of sound position. The random error <u>*s*</u> for each participant was computed as the standard deviation of the responses at each sound position, averaged across all sound. Figure 9B shows change in 3D-error in the two listening conditions. Considering all participants and trials, the improvement in sound localization in active (28.5 ± 2.4) compared to static listening (32.2 ± 2.3) emerged as marginally significant (F(1, 19) = 4.34, p = .051, 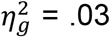). However, when the difference was studied as a function of mean number of head movements a positive correlation emerged (r = 0.36, p = 0.027). The higher the proportion of trials with head movements during sound emission, the larger the performance improvement in active compared to static listening. A convergent correlation emerged also between mean number of head-movements and 3D-error (r = 0.34, p = 0.038).

**Figure 9.**
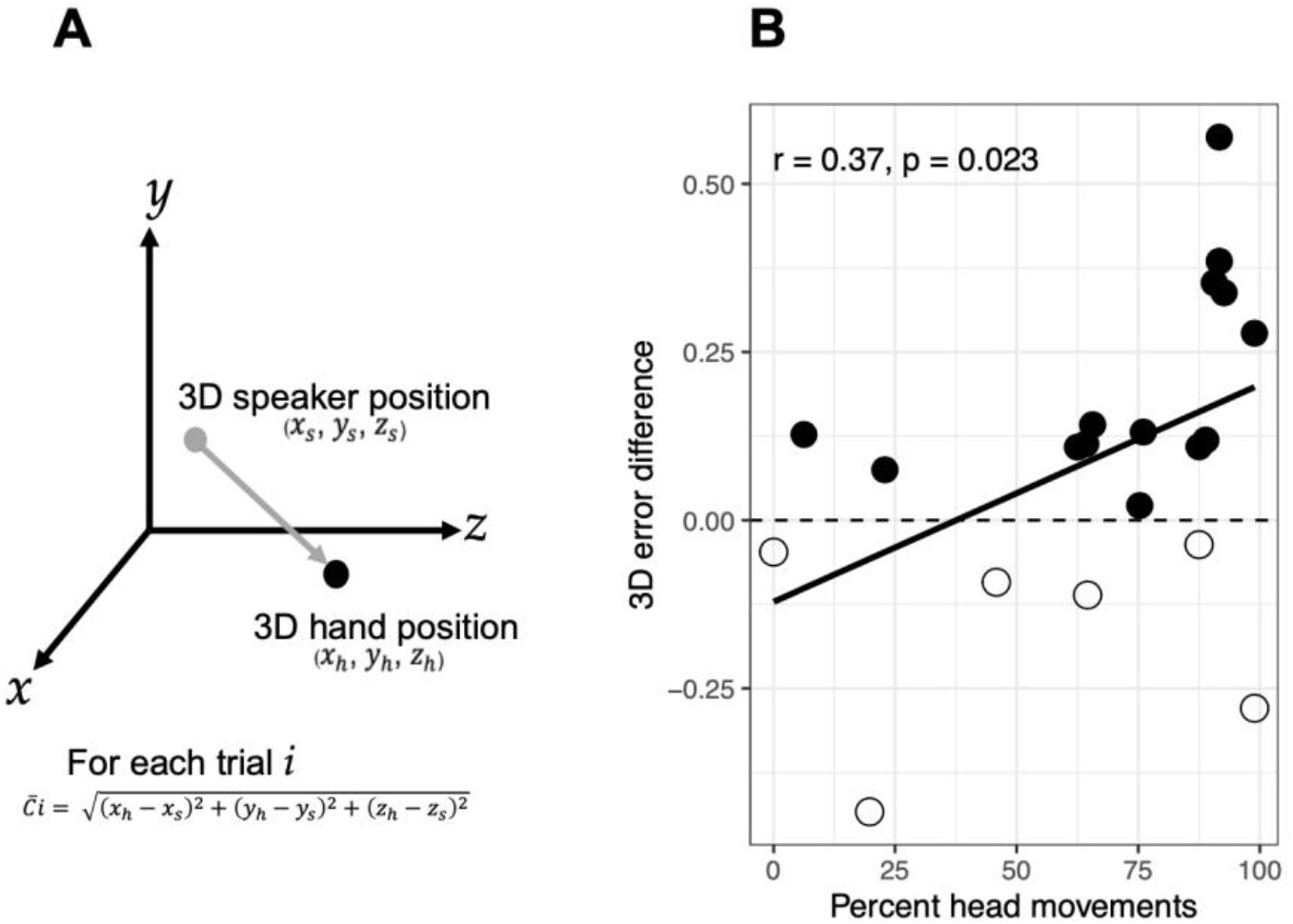
3D-error. (A) Schematics for the calculation of the 3D vector. (B) Scatterplot of the difference in 3D-error between active and static listening (normalized difference based on static listening performance), as a function of percent head movements. Filled circles indicate participants who improved in active compared to static listening; empty circles indicate participants whose sound localization performance decreased in active compared to static listening.

## DISCUSSION

In the present study, we aimed to report on a newly developed system for measuring sound localization, that allows replicable delivery of real sounds in 3D with respect to the head, while leaving participants free in their head-movement behavior. To validate our system, we examined to what extent voluntary head-movements improved sound localization in 3D – i.e., azimuth, elevation, and depth – by comparing static vs. active listening postures. We showed that our system is effective for controlling the delivery of sounds in 3D while measuring in real-time the participant’s behavior (hand, head and eye position), without imposing a physically constrained posture and with minimal training required. In addition, we found that head-behavior changes sound localization in 3D, improving specifically azimuth and elevation perception.

### Sound localization in 3D

Despite minimal physical constraints, we succeeded in presenting free-field sounds at controlled 3D locations, across trials, participants, and sessions led by different experimenters. When referenced to the center of the head, the average error across target positions for all participants and sessions of recording was below 1 cm. Notably, all experimenters achieved accurate and fast speaker 3D positioning with a few minutes training (< 5 minutes). Most importantly, participants required no procedural training to perform the task, with less than 7% of trials rejected for no-compliance to instructions.

Previous studies that examined sound localization in 3D were based on one of two approaches. On the one hand, some studies have used several real speakers in external space (free field sounds), which occupied fixed locations with respect to the participant (Ahrens et al., 2019), or a single speaker, which is moved and tracked in space (Brungart et al., 1999). On the other hand, other studies have used virtual auditory stimuli, either obtained by recording real sounds from the ear-canal of individual participants, or generated synthetically using Head-Related Transfer Functions HRTFs (individualized or not).

In the experimental approaches based on free field sounds, it is practically mandatory that the participant keeps a fixed posture, because this is the only way to establish off-line the position of target sounds with respect to the head (Wightman & Kistler 1992; Oldfield & Parker 1984; Brungart et al., 1999; Seeber et al., 2004). In addition, because physical sound sources are present, these approaches face the problem of controlling the contribution of visual cues to sound localization, as any visual prior will contribute to the interpretation of auditory cues (Jackson & Morton 1984; Makous & Middlebrooks 1990). Participants are therefore blind-folded, sometimes from the moment they enter the experimental room (e.g., Ahrens and al., 2019), or are instructed to close their eyes at specific moments during the task (e.g., Brungart et al., 1999), or face speakers hidden behind a fabric panel of some sort (e.g., Rabini et al., 2019).

Although studies using this approach typically used speakers at fixed positions in the environment, this needs not be the case, as first proposed by Brungart and colleagues (Brungart et al., 1999). In their pioneering study, participants were required to indicate the position of the sound source around their head while the experimenter manually placed the speaker during each trial. To ensure reproducibility of sound source coordinates across trials and participants, the participant’s head was immobilized with a chin rest and the experimenter received verbal instructions about the pre-determined speaker location (i.e., a set of three numbers spoken through headphones, from 1 to 6, to indicate azimuth, elevation and distance according to learned correspondences). Although the speaker position was somewhat approximate, its actual location was recorded at the end of each trial using a position-sensing system mounted on the chin rest. Using this real sound method, Brungart and colleagues (1999) succeeded in placing sound sources in 3D space. While innovative, this experimental setup was complex and time consuming and, unfortunately, it was not followed-up by other investigators Participants had to familiarize with the procedure before data collection.

Moreover, the method had intrinsic limitations for the study of sound perception. First, sound source positions were variable among participants because the speaker’s coordinates were interpreted by the experimenter in each trial using a number-to-coordinate mapping. Second, subjects had to close their eyes during sound positioning, thus limiting most oculomotor information that could have enhanced sound localization abilities (Maddox et al., 2014) and preventing any eye-movement monitoring.

With respect to the pioneering approach proposed by Brungart and colleagues (Brungart et al., 1999), our novel methodology allows: (1) accurate guiding of the speaker to pre-determined coordinates, that need not be interpreted by the experimenter; (2) anchoring of all coordinates to head position, without using a chinrest; (3) control of visible cues about speaker position; and (4) control of eye position throughout the experiment. In addition, with respect to more classic approaches based on fixed speakers, our paradigm allowed sampling positions all around the participant in 3D. This is particularly evident when studying distance: if fixed speakers are used, placing them at different distances would interfere with the sound field generated by one another.

In the experimental approaches based on virtual auditory stimuli, two alternative strategies have been employed: in-ear sounds encoding and synthetic sound encoding. In in-ear sounds encoding, real sounds originating from pre-determined locations are recorded from the ear-canal of individual participants or from microphones in a dummy-head. The head has to remain motionless because it is impossible to modify the sound scene after having encoded each sound. In synthetic sound encoding, sound positions are computed in post-production via specialization techniques (HRTFs or Binaural Room Impulse Response) including several cues related to body features or room. Although a considerable amount of fundamental knowledge about HRTFs has accumulated over the years (for review, see Lida, 2019), synthetic sound encoding is not as powerful as the approach based on real speakers because several features must be implemented in the delivered sound.

First, the HRTFs depends on each individual’s morphology (size of the head, shape of the pinna, etc.) which varies from one listener to another. The use of HRTFs without taking into account the individual characteristics of the subject’s head (non-individualized HRTF), leads to the occurrence of front-back confusions, distortions of spatial and timbre perception (Seeber & Fastl, 2003; Wenzel et al., 1993; Middlebrooks, 1999), and synthetic sounds sources delivered by ear-phones are perceived inside the head, thus without distance perception (Begault et al., 2001). Second, modelling sound space with HRTFs is still a major challenge, particularly in relation to distance perception. Modelling sound distance requires a combination of a variety of acoustic cues (for review, see Zahorik et al., 2005): intensity of sound, direct-to-reverberant energy ratio, frequency spectrum, binaural cues. In addition, non-acoustic visual cues are often added to improve distance perception. To add to the complexity, different HRTFs models may be necessary when dealing with acoustic sources in far vs. near acoustic spaces (Brungart & Rabinowitz 1999). This is probably the reason why few of synthetic sound studies have dealt with sound localization in the 3D space reachable by the hand and close to the head. More generally, it may explain why distance is implemented separately with respect to the other dimensions: changes in azimuth and elevation are typically implemented at fixed distances (e.g., Rajendran & Gamper 2019; Lubeck et al., 2019), whereas changes in distance are typically implemented with fixed azimuth and elevation (e.g., Zahorik, 2002); (3) Third, use of synthetic sounds becomes extremely problematic when dealing with people with hearing deficit, individuals using hearing aids or cochlear implants, and children -- for whom HRTFs are likely to change faster over time. Yet, notable attempts to study these populations with HRTFs exist in the literature (e.g., Majdak et al., 2011, for cochlear implants users). In the Majdak and colleagues study (2011), they delivered synthetic sounds directly to hearing aids of participants. To allow comparable sound perception across cochlear implant participant, the levels of synthetic sounds were adapted on an individual basis. However, despite this procedure, the authors faced the problem of variability of loudness perception over sessions. Finally, when compared to real-sound stimulation, the authors pointed that their “artificial testing conditions overestimated the real-life localization performance of our listener” (Majdak et al., 2011, page 207).

Several aspects of our approach to sound localization are notable with respect to the approach based on virtual auditory stimuli: (1) our procedure based on real sounds does not require measuring individual HRTF or recording from the individual’s ear canal; (2) the three dimensions can easily be changed at the same time in the whole space, particularly in reaching space; (3) our 3D sound localization approach can easily be adopted in individuals who use hearing aids or cochlear implants, children and elderly, for which synthetic acoustic approach is possible yet extremely time consuming and challenging.

Using the SPHERE system to characterize sound localization performance of normal hearing subject in static listening posture, we were able to show that participants discriminated azimuth, elevation and distance. Absolute average errors were 21.6 degrees, 25.1 degrees and 10.5 cm respectively. Differences emerged across along the medial dimension, with poorer performance in azimuth for front (26.1 degrees) compared to back targets (17.1 degrees), which emerged as a bias to point to more eccentric positions for frontal sources. For elevation, inaccuracy appeared for frontal sources compared to back ones (even if did not reach significant difference). This overall angular error measured in our study is numerically greater than those obtained by others studies in free field sound stimulation. For comparable elevation position, Brungart and colleagues (1999) obtained a mean angular error of 16.3 degrees, and Wightman & Kistler (1999), a mean error of 21.1 degrees. While the performance of our normal-hearing adults may appear poor, three aspects of our paradigm may contribute to this outcome. First, participants were only informed that target sounds would be delivered within reaching distance, but had no further information on their positions -- i.e., they expected sounds to appear all around the body. Second, these errors combine uncertainty across the third dimension (i.e. distance is also unknown for participants). Third, they had no visual prior that could help sound localization.

### Head-movements

Space is not represented directly on inner ear and spatial cues must be derived by the nervous system from incoming sound and dynamic auditory cues (i.e. head motion). Head motion is spontaneous in natural listening (Kim et al., 2013), and its key role in sound localization has been proposed since the last century (Wallach, 1940; see also Wightman & Kistler, 1999). It is generally assumed that head movements are accounted for during the computation of sound-source coordinates (Goossens & Van Opstal, 1999), as suggested by the fact that neck muscle stimulation has been shown to produce a shift of auditory localization toward the side of stimulation (Lewald et al., 1999). Converging evidence suggests that continuous integration of head motion feedback on auditory input provides more stable sound source position (Vliegen et al., 2004). Several psychoacoustic studies demonstrate that head motion increases sound localization in humans (Pollack & Rose 1967; Perrett & Noble, 1997a; 1997b; Wightman & Kistler 1999; Vliegen et al., 2004; Brimijoin at al., 2013; Honda et al., 2013; McAnally & Martin, 2014), and monkeys (Populin, 2006). Speech perception improvements have also been documented (Munhall et al., 2004), even on patients with hearing aids (Mueller et al., 2014). Finally, head movements also proved important for the experience the surrounding ambient sound (Suzuki et al., 2011) and are starting to be implemented in machine-hearing systems (Ma et al., 2015).

Notwithstanding these shared considerations, the study of head-movements in sound localization remained largely overlooked. Searching with the descriptor ‘sound localization’ in the MEDLINE repository since the work of Wallace (1940) returns 6014 entries, but when including the additional descriptor ‘head-movement’ reduces the entries to 271 articles related to spatial hearing performance (4.5%). One reason for this discrepancy may reside in the fact that taking into account head-movement has been problematic for most approaches to sound localization. In free-field approaches, head-movements have mostly been prevented, to ensure reproducibility of sound source position across trials and participants, or remained uncontrolled. In in-ear approaches, head tracking enables the signals fed to the two ears to change accordingly to head movements, so that the perception of the virtual sound source remains while moving the head. Indeed, some psychoacoustic studies have implemented in their model a dynamic modification of HRTFs cues caused by head motion (Brimijoin et al., 2013; Hendrickx et al., 2017; Pöntynen & Salminen 2019). Motion allows listeners to take advantage of dynamic localization cues, i.e. the changes in acoustic input caused by the movement of the head with respect to the sound source. This dynamic change caused by head movements can enhance sound perception: the accuracy of sound localization is better than that when HRTFs are not switched. Despite advances in 3D sound modelling taking particular account of head motion, mostly HRTF applications did not use 3D head tracking displacement (i.e. rotation in azimuth, tipping and tilting) and limit HRTF updating to lateral head movements only (Ma et al., 2015).

Our novel approach overcomes previous limitations by measuring head-position in real-time, thus allowing accurate and reproducible positioning of sounds even without physical constraints to head-position, and a full description of head behavior before, during and after sound presentation. Notably, in the active listening posture, no specific head-movement strategy was imposed on participants, and they were free to move or not their head to explore their surrounding environment during sound emission. As a proof-of-concept of our methodology, we chose to compare static vs. active listening posture for sounds lasting 3 seconds, a duration for which participants can benefit of acoustic dynamic cues from head-motion (Pollack & Rose, 1967; review by Middlebrooks & Green 1991).

### Active listening improves 3D sound localization

We found that head motion during sound emission improved both their azimuth and elevation performances. Improvements concerned azimuth and elevation of front sources localizations (both absolute and variable errors) and azimuth and elevation of back sources localizations (variable error). For the third dimension (depth), active listening did not modify performances.

Previous studies have shown that head movements help normal hearing listeners to distinguish between sounds coming from front and rear positions (Makous & Middlebrooks 1990; Mueller et al., 2014; Bronkhorst 1995; Mackensen, 2004; Perrett & Noble 1997a; Wallach 1940; Wenzel et al., 1993; Wightman & Kistler, 1999). Front-back confusion is removed if head movements were larger than 5° (Perrett & Noble, 1997a), and if sounds were long enough (Muller et al., 2014; Perrett & Noble, 1997a). For a 3-sec. sound duration, in Perret and Noble’s study (1997a), a few confusions occurred but only in situations where the listeners did not rotate their heads. In our study with a sound duration of 3 sec., the subjects had enough time to improve their performance in localizing sound sources by head motion.

### Current limitation of our approach

Sound localization studies with head motionless reported a more accurate performance for sources located in the frontal space than in back space (e.g. Oldfield & Parker, 1984; Brungart et al., 1999). In the present study, when head motion was not allowed, participants were more accurate in localizing sounds presented in the back (17.1 degrees) compared to front space (26.1 degrees). However, their performance increased when head motion was allowed (back: 16.7 degrees; front: 16.0 degrees). There are two likely reasons for this finding. First, several studies (Etchemendy et al., 2018; Brungart et al., 2000; Haber et al., 1993) showed superior accuracy and lower variability with direct-location response methods for sound localization (e.g., pointing with body parts) and also helps front-back discrimination (Aggius-Vella and al., 2017). In Oldfield and Parker study (1984), participant were head fixed and used a gun to localize the sound direction. The error in azimuth localization increased in the regions behind the head, particularly for azimuth positions 120° to 160°. When pointing at sounds behind the head subjects were instructed to turn the gun backwards and to ‘shoot’ from the front through their head to the sound source. In our direct-method, subjects are free to orient their body and their arm towards the rear field to respond, leading to decrease errors in the back space. Here we add to this direct-location panoply of methods an ecological, sensitive and rather intuitive (see Valzolgher et al., under revision) direct-location response to study 3D human spatial audition in the reaching space. Second, it has been reported that wearing and HMD may alter sound localization cues (Gupta et al., 2018; Ahrens et al., 2019; Genovese et al., 2018). Here we add that it may also affect sound source localization accuracy. In the Ahrens and colleagues study (2019), which explored the effect of HMD on sound localization presented in a frontal plane, a larger azimuthal error was found for lateral sound sources when participants whore HMD than without. In this study, sound duration was very short (240 ms) to limit the effect of head movement during sound presentation. These findings are in keeping with those observed here in static listening conditions (i.e., head motionless) for the frontal sources and could be due to a binaural disparity caused by the HMD. This possibility is supported by the participants azimuthal and elevation improvement for frontal targets during the active listening condition. Accordingly, the lateral error increase due to HMD does not exist for the rear sources in either static or active listening, as the HMD is no longer offer an obstacle for binaural processing of rear sounds. In regard to depth, participants succeed in distinguish the three distances sound sources with an underestimation for far targets compared to near ones during stating listening. The commonness of errors in distance localization is well-established (Brungart, 1999; Kearney et al., 2010; Zahorik 2002; 2005; Zahorik & Wightman 2001; Middlebrooks & Green, 1991; Parseihian et al., 2014). Sound distance cues such as binaural cue and interaural level difference (Brungart, 1999) are important for distance localization in the near-head acoustic field and not affected by the HMD.

## CONCLUSION

SPHERE (European patent n° WO2017203028A1) is a valid tool to accurately sample the spatial abilities in auditory perception all around the subject, with minimal constraints to participant and experimenter. Most interestingly, SPHERE proved sensitive to allow detecting and quantifying the contribution of active listening (here, free head motion during sound emission) to improve sound localization accuracy and precision. This system also paves the way for future research, clinical and industrial applications that will leverage the full potential offered by having embedded a VR HMD in the SPHERE system. Indeed, sound localization is not only a purely acoustic phenomenon, but a combination of multisensory information processing, especially relying on visual environmental information. HMDs offer the possibility of a 3D, fully immersive visual experience that combined to 3D sound simulation and head-body motion tracking will offer a highly versatile opportunity to assess normal and pathological sound localization performance. Beyond the range of performance of normal hearing people, it is a tool of choice for patients with cochlear implant and / or hearing aids (Avan et al. 2015).

## Supporting information

Supplementary Informations GAVEAU et al

## Acknowledgements

This work was supported by LabEx CORTEX (https://labex-cortex.universite-lyon.fr), the Medisite Fundation (https://www.fondationdefrance.org/fr/fondation/fondation-medisite), ITMO New technologies for Neuroscience, Idex Lyon ‘Senses In Space Lab’, ANR (ANR-16-CE17-0016-01), CNRS-PICS (Programme PHC Galilée Univ Franco-Italienne), and from IHU CeSaMe ANR-10-IBHU-0003.

## REFERENCES

Aggius-Vella, E., Campus, C., Finocchietti, S., & Gori, M. (2017). Audio Spatial Representation Around the Body. Frontiers in Psychology, 8, 1932. https://doi.org/10.3389/fpsyg.2017.01932

Ahrens, A., Kasper Duemose, L., Marschall, M., & Dau, T. (2019). Sound Source Localization with Varying Amount of Visual Information in Virtual Reality. PLoS ONE 29 (14): 1. https://doi.org/10.1371/journal.pone.0214603

Andéol, G., Savel, S., & Guillaume, A. (2014). Perceptual factors contribute more than acoustical factors to sound localization abilities with virtual sources. Front Neurosci. 2014; 8: 451. doi: 10.3389/fnins.2014.00451

Andéol, G., & Simpson, B. D. (2016). Editorial: How, and Why, Does Spatial-Hearing Ability Differ among Listeners? What is the Role of Learning and Multisensory Interactions? Frontiers in Neuroscience, 10:36. https://doi.org/10.3389/fnins.2016.

Avan, P., Giraudet, F., & Büki, B. (2015). Importance of Binaural Hearing ». Audiology and Neurotology 20 (1): 3–6. https://doi.org/10.1159/000380741

Bahill, A.T., & McDonald J.D. (1983). Frequency limitations and optimal step size for the two-point central difference derivative algorithm with applications to human eye movement data. IEEE Trans Biomed Eng. 30(3):191–4. doi: 10.1109/tbme.1983.325108

Begault, D.R., Wenzel, E.M., & Anderson, M.R. (2001). Direct Comparison of the Impact of Head Tracking, Reverberation, and Individualized Head-Related Transfer Functions on the Spatial Perception of a Virtual Speech Source. J. Audio Eng. Soc. 49: 14.

Benjamini, Y., & Hochberg, Y. (1995). Controlling the false discovery rate: a practical and powerful approach to multiple testing. Journal of the Royal Statistical Society, Series B (Methodological). 57(1): 289–300.

Brimijoin, W.O., McShefferty, D., & Akeroyd. M.A. (2010). Auditory and Visual Orienting Responses in Listeners with and without Hearing-Impairment. The Journal of the Acoustical Society of America 127 (6): 3678–88. https://doi.org/10.1121/1.3409488.

Brimijoin, W.O., McShefferty, D., & Akeroyd, M.A. (2012). Undirected Head Movements of Listeners with Asymmetrical Hearing Impairment during a Speech-in-Noise Task. Hearing Research 283 (1-2): 162–68. https://doi.org/10.1016/j.heares.2011.10.009.

Brimijoin, W.O., Boyd, A.W., & Akeroyd, M.A. (2013). The Contribution of Head Movement to the Externalization and Internalization of Sounds. PLoS ONE 8 (12): e83068. https://doi.org/10.1371/journal.pone.0083068.

Bronkhorst, A.W. (1995). Localization of Real and Virtual Sound Sources ». The Journal of the Acoustical Society of America 98 (5): 2542–53. https://doi.org/10.1121/1.413219.

Brungart, D.S. (1999). Auditory localization of nearby sources. III. Stimulus effects. J Acoust Soc Am. 1999 Dec;106(6):3589–602. https://doi.org/10.1121/1.428212

Brungart, D.S., Durlach, N.I., & Rabinowitz., W.M. (1999). Auditory Localization of Nearby Sources. II. Localization of a Broadband Source. The Journal of the Acoustical Society of America 106 (4): 1956–68. https://doi.org/10.1121/1.427943.

Brungart, D.S., & Rabinowitz., W.M. (1999). Auditory Localization of Nearby Sources. Head-Related Transfer Functions. The Journal of the Acoustical Society of America 106 (3): 1465–79. https://doi.org/10.1121/1.427180.

Brungart, D.S., Rabinowitz, W.M., & Durlach, N.I. (2000). Evaluation of Response Methods for the Localization of Nearby Objects. Perception & Psychophysics 62 (1): 48–65. https://doi.org/10.3758/BF03212060.

Bulkin, D.A, & Groh, J.M. (2006). Seeing Sounds: Visual and Auditory Interactions in the Brain. Current Opinion in Neurobiology 16 (4): 415–19. https://doi.org/10.1016/j.conb.2006.06.008.

Etchemendy, P.E., Spiousas, I., Calcagno, E.R., Abregú, E., Eguia, M.C., & Vergara, R.O. (2018). Direct-Location versus Verbal Report Methods for Measuring Auditory Distance Perception in the Far Field. Behavior Research Methods 50 (3): 1234–47. https://doi.org/10.3758/s13428-017-0939-x.

Genovese, A., Zalles, G., Reardon, G., & Roginska, A. Acoustic perturbations in hrtfs measured on mixed reality headsets. In Audio Engineering Society Conference: 2018 AES International Conference on Audio for Virtual and Augmented Reality. Audio Engineering Society.

Goossens, H.H.L.M., &. van Opstal, A.J. (1999). Influence of Head Position on the Spatial Representation of Acoustic Targets. Journal of Neurophysiology 81 (6): 2720–36. https://doi.org/10.1152/jn.1999.81.6.2720.

Grantham, D.W., Ashmead, D.H., Ricketts, T.A., Labadie, R.F., & Haynes, D.S. (2007). Horizontal-Plane Localization of Noise and Speech Signals by Postlingually Deafened Adults Fitted With Bilateral Cochlear Implants. Ear and Hearing 28 (4): 524–41. https://doi.org/10.1097/AUD.0b013e31806dc21a.

Grantham, D.W., Ricketts, T.A., Ashmead, D.H., Labadie, R.F., & Haynes, D.D. (2008). Localization by postlingually deafned adults fittes with a single cochlear implant. The Larynoscope, 118(1), 145–151. https://doi.org/10.1097/MLG.0b013e31815661f9

Groh, J.M., & Sparks, D. L. (1992). Two Models for Transforming Auditory Signals from Head-Centered to Eye-Centered Coordinates ». Biological Cybernetics 67 (4): 291–302. https://doi.org/10.1007/BF02414885.

Gupta, A., Ranjan, R., He, J., & Gan, W-S. (2018). Investigation of Effect of VR/AR Headgear on Head Related Transfer Functions for Natural Listening. AES International Conference on Audio for Virtual and Augmented Reality (August 2018). http://www.aes.org/e-lib/browse.cfm?elib=19697

Haber, L., Haber, R.N., Penningroth, S., Novak, K., & Radgowski, H. (1993). Comparison of Nine Methods of Indicating the Direction to Objects: Data from Blind Adults. Perception 22 (1): 35–47. https://doi.org/10.1068/p220035.

Hendrickx, E., Stitt, P., Messonnier, J-C., Lyzwa, J-M., Katz, B.F.G, & de Boishéraud, C. (2017). Influence of Head Tracking on the Externalization of Speech Stimuli for Non-Individualized Binaural Synthesis. The Journal of the Acoustical Society of America 141 (3): 2011–23. https://doi.org/10.1121/1.4978612.

van Hoesel, R.J.M., & Tyler, R.S. (2003). Speech Perception, Localization, and Lateralization with Bilateral Cochlear Implants. The Journal of the Acoustical Society of America 113 (3): 1617–30. https://doi.org/10.1121/1.1539520.

Honda, A., Shibata, H., Hidaka, S., Gyoba, J., Iwaya, Y., & Suzuki, Y. (2013). Effects of Head Movement and Proprioceptive Feedback in Training of Sound Localization. I-Perception 4 (4): 253–64. https://doi.org/10.1068/i0522.

Jackson, A., Morton, J. (1984). Facilitation of Auditory Word Recognition. Memory & Cognition 12 (6): 568–74. https://doi.org/10.3758/BF03213345.

Kearney, G., Gorzel, M., Boland, F., & Rice, H., (2010). Depth perception in interactive virtual acoustic environments using higher order ambisonic soundfields. Proc. of the 2nd International Symposium on Ambisonics and Spherical Acoustics.

Kim, C., Mason, R., & Brookes, T. (2013). Head Movements Made by Listeners in Experimental and Real-Life Listening Activities. Journal of the Audio Engineering Society, 2013, sect. 61(6). http://www.aes.org/e-lib/browse.cfm?elib=16833

Kumpik, D.P., Campbell, C., Schnupp, J.W.H., & King, A.J. (2019). Re-weighting of sound localization cues by audiovisual training. Frontiers in Neuroscience, 13: 1164. https://doi.org/10.3389/fnins.2019.01164

Lewald, J., & Ehrenstein, W.H. (1996). The Effect of Eye Position on Auditory Lateralization. Experimental Brain Research, no 108: 473–85. dio 10.1007/bf00227270

Lewald, J., Karnath, H-O., & Ehrenstein, W.H. (1999). Neck-Proprioceptive Influence on Auditory Lateralization. Experimental Brain Research 125 (4): 389–96. https://doi.org/10.1007/s002210050695.

Lida, K. (2019). Head-Related Transfer Functions And Acoustic Virtual Reality. Eds Springer Nature. ISBN 9789811397455

Litovsky, R., Parkinson, A., & Arcaroli, J. (2009). Spatial Hearing and Speech Intelligibility in Bilateral Cochlear Implant Users. Ear and Hearing 30 (4): 419–31. https://doi.org/10.1097/AUD.0b013e3181a165be.

Lubeck, T., Arend, J.M., & Porschmann, C. (2019). HMD-Based Virtual Environments for Localization Experiments. Conference DAGA 2019 Rostock, pp. 1116–1119.

Ma, N., May, T., Wierstorf, H., & Brown, G.J. (2015). A Machine-Hearing System Exploiting Head Movements for Binaural Sound Localisation in Reverberant Conditions. In 2015 IEEE International Conference on Acoustics, Speech and Signal Processing (ICASSP), 2699–2703. South Brisbane, Queensland, Australia: IEEE. https://doi.org/10.1109/ICASSP.2015.7178461.

Mackensen, P. (2004). Auditive Localization. Head Movements, an Additional Cue in Localization. PhD Thesis, Berlin: Technische Universität Berlin.

Maddox, R.K., Pospisil, D.A., Stecker, G.C., & Lee, A.K.C. (2014). Directing Eye Gaze Enhances Auditory Spatial Cue Discrimination. Current Biology 24 (7): 748–52. https://doi.org/10.1016/j.cub.2014.02.021.

Majdak, P., Goupell, M.J. & Laback, B. (2011). Two-Dimensional Localization of Virtual Sound Sources in Cochlear-Implant Listeners. Ear and Hearing 32 (2): 198–208. https://doi.org/10.1097/AUD.0b013e3181f4dfe9.

Makous, J.C., & Middlebrooks, J.C. (1990). Two-dimensional Sound Localization by Human Listeners. The Journal of the Acoustical Society of America 87 (5): 2188–2200. https://doi.org/10.1121/1.399186.

McAnally, K.I., & Martin, R.L. (2014). Sound Localization with Head Movement: Implications for 3-d Audio Displays. Frontiers in Neuroscience 8:210. https://doi.org/10.3389/fnins.2014.00210.

Middlebrooks, J.C, & Green, D.M. (1991). Sound Localization by Human Listeners. Annu Rev Psychol 42: 135–59. doi: 10.1146/annurev.ps.42.020191.001031

Middlebrooks, J.C. (1999). Individual Differences in External-Ear Transfer Functions Reduced by Scaling in Frequency. The Journal of the Acoustical Society of America 106 (3): 1480–92. https://doi.org/10.1121/1.427176.

Mueller, M.F, Meisenbacher, K., Lai W-K., & Dillier, N. (2014). Sound Localization with Bilateral Cochlear Implants in Noise: How Much Do Head Movements Contribute to Localization? Cochlear Implants International 15 (1): 36–42. https://doi.org/10.1179/1754762813Y.0000000040.

Munhall, K.G., Jeffery J.A., Callan, D.E., Kuratate, T., & Vatikiotis-Bateson, E. (2004). Visual Prosody and Speech Intelligibility: Head Movement Improves Auditory Speech Perception. Psychological Science 15 (2): 133–37. https://doi.org/10.1111/j.0963-7214.2004.01502010.x.

Nava, E., Bottari, D., Bonfioli, F., Beltrame, M.A., & Pavani, F. (2009). Spatial Hearing with a Single Cochlear Implant in Late-Implanted Adults. Hearing Research 255 (1-2): 91–98. https://doi.org/10.1016/j.heares.2009.06.007.

Oldfield, S.R, & Parker, S.P.A. (1984). Acuity of Sound Localisation: A Topography of Auditory Space. I. Normal Hearing Conditions. Perception 13 (5): 581–600. https://doi.org/10.1068/p130581.

Parseihian, G., Jouffrais, C., Katz, B.F.G. (2014). «Reaching Nearby Sources: Comparison between Real and Virtual Sound and Visual Targets ». Frontiers in Neuroscience 8: 269. https://doi.org/10.3389/fnins.2014.00269.

Pavani, F., Meneghello F., Làdavas E. (2001). Deficit of auditory space perception in patients with visuospatial neglect. Neuropsychologia. 39(13):1401–9. Doi. 10.1016/s0028-3932(01)00060-4

Pavani, F., Farnè, A., & Làdavas, E. (2003). Task-Dependent Visual Coding of Sound Position in Visuospatial Neglect Patients. NeuroReport 14 (1) 99–103. https://doi.org/10.1097/00001756-200301200-00019.

Pavani, F., Husain, M., & Driver, J. (2008). Eye-Movements Intervening between Two Successive Sounds Disrupt Comparisons of Auditory Location. Experimental Brain Research 189 (4): 435–49. https://doi.org/10.1007/s00221-008-1440-7.

Perrett, S., & Noble, W. (1997a). The Contribution of Head Motion Cues to Localization of Low-Pass Noise. Perception & Psychophysics 59 (7): 1018–26. https://doi.org/10.3758/BF03205517.

Perrett, S., & Noble, W. (1997b). The Effect of Head Rotations on Vertical Plane Sound Localization. The Journal of the Acoustical Society of America 102 (4): 2325–32. https://doi.org/10.1121/1.419642.

Pollack, I., & Rose, M. (1967). Effect of Head Movement on the Localization of Sounds in the Equatorial Plane. Perception & Psychophysics 2 (12): 591–96. https://doi.org/10.3758/BF03210274.

Pöntynen, H., & Salminen, N.H. (2019). Resolving Front-Back Ambiguity with Head Rotation: The Role of Level Dynamics. Hearing Research 377: 196–207. https://doi.org/10.1016/j.heares.2019.03.020.

Populin, L.C. (2006). Monkey Sound Localization: Head-Restrained versus Head-Unrestrained Orienting. Journal of Neuroscience 26 (38): 9820–32. https://doi.org/10.1523/JNEUROSCI.3061-06.2006.

Rabini, G., Altobelli, E., Pavani, F. (2019). Interactions between egocentric and allocentric spatial coding of sounds revealed by a multisensory learning paradigm. Sci Rep. 2019; 9: 7892. doi: 10.1038/s41598-019-44267-3

Rajendran, V.G., & Gamper, H. (2019). Spectral Manipulation Improves Elevation Perception with Non-Individualized Head-Related Transfer Functions. The Journal of the Acoustical Society of America 145 (3): EL222–28. https://doi.org/10.1121/1.5093641.

Rakerd, B., & Hartmann, W.M. (1986). Localization of Sound in Rooms, III: Onset and Duration Effects. The Journal of the Acoustical Society of America 80 (6): 1695–1706. https://doi.org/10.1121/1.394282.

Seeber, B.U., & Fastl, H. (2003). Subjective Selection of Non-Individual Head-Related Transfer Functions. Proceedings of the 2003 International Conference on Auditory Display, Boston, MA, USA, July 6–9. http://hdl.handle.net/1853/50488

Seeber, B.U., Baumann, U., & Fastl, H. (2004). Localization Ability with Bimodal Hearing Aids and Bilateral Cochlear Implants. The Journal of the Acoustical Society of America 116 (3): 1698–1709. https://doi.org/10.1121/1.1776192.

Slattery, W.H., & Middlebrooks, J.C. (1994). Monaural Sound Localization: Acute versus Chronic Unilateral Impairment I ». Hearing Research 75(1-2): 38–46. doi. 10.1016/0378-5955(94)90053-1

Suzuki, Y., Brungart, D., Iwaya, Y., Iida, K., Cabrera, D., & Kato. H. (2011). Principles and Applications of Spatial Hearing. Eds. World Scientific. https://doi.org/10.1142/7674.

Távora-Vieira, D., De Ceulaer, G., Govaerts, P.J., & Rajan, G.P. (2015). Cochlear Implantation Improves Localization Ability in Patients With Unilateral Deafness. Ear and Hearing 36 (3): e93–98. https://doi.org/10.1097/AUD.0000000000000130.

Valzolgher, C., Verdelet, G., Salemme, R., Lombardi, L., Farnè, A., Pavani, P. Reaching to sounds in virtual reality: A multisensory-motor approach to re-learn sound localisation. Under review

Vliegen, J., van Grootel, T.J., & van Opstal, A. J. (2004). Dynamic Sound Localization during Rapid Eye-Head Gaze Shifts ». Journal of Neuroscience 24 (42): 9291–9302. https://doi.org/10.1523/JNEUROSCI.2671-04.2004.

Wallach, H. (1940). The Role of Head Movements and Vestibular and Visual Cues in Sound Localization. Journal of Experimental Psychology 27 (4): 339–68. https://doi.org/10.1037/h0054629.

Wenzel, E.M., Arruda, M., Kistler, D.J., & Wightman, F.L. (1993). Localization Using Nonindividualized Head-related Transfer Functions ». The Journal of the Acoustical Society of America 94 (1): 111–23. https://doi.org/10.1121/1.407089.

Wightman, F.L., & Kistler, D.J. (1992). The Dominant Role of Low-frequency Interaural Time Differences in Sound Localization ». The Journal of the Acoustical Society of America 91 (3): 1648–61. https://doi.org/10.1121/1.402445.

Wightman, F.L., & Kistler, D.J. (1999). Resolution of Front–Back Ambiguity in Spatial Hearing by Listener and Source Movement. The Journal of the Acoustical Society of America 105 (5): 2841–53. https://doi.org/10.1121/1.426899.

Zahorik, P, & Wightman, F.L. (2001). Loudness Constancy with Varying Sound Source Distance. Nature Neuroscience 4 (1): 78–83. https://doi.org/10.1038/82931.

Zahorik, P. 2002. Assessing Auditory Distance Perception Using Virtual Acoustics. The Journal of the Acoustical Society of America 111 (4): 1832–46. https://doi.org/10.1121/1.1458027.

Zahorik, P., Brungart, D.S., Bronkhorst, A.W. (2005). Auditory Distance Perception in Humans: A Summary of Past and Present Research. Acat acustica United with Acustica. vol. 91(3): 409–420

