## Supplementary Informations GAVEAU et al for "SPHERE: A novel approach to 3D and active sound localization"

### Supplementary figures

Gaveau et al.

### Figure S1

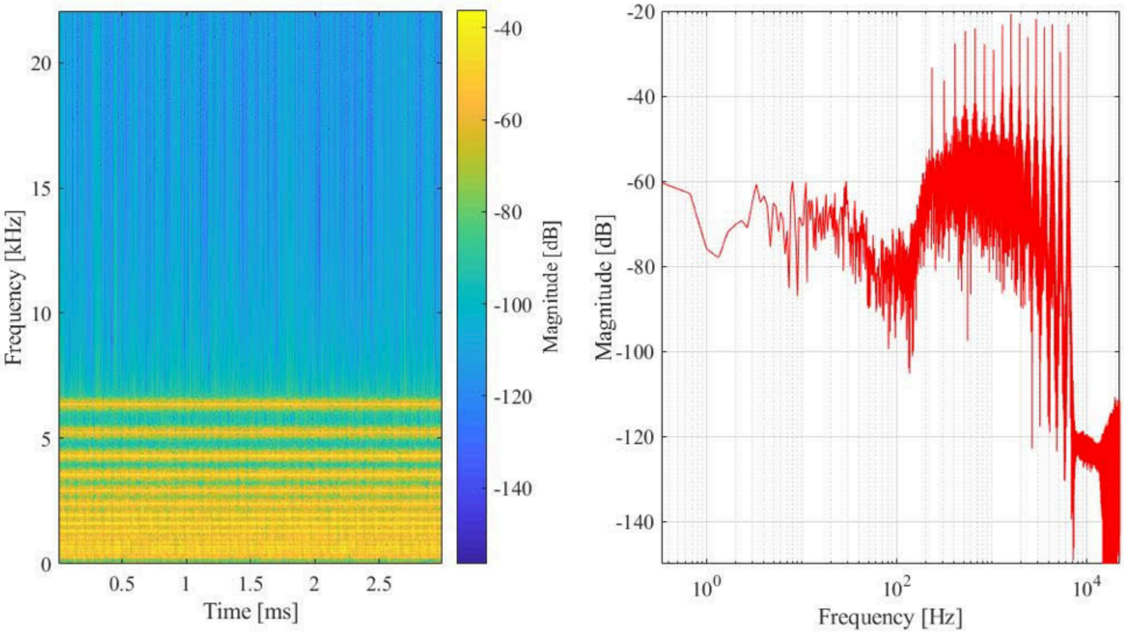


**Figure S1.** Spectrograms as a function of time and magnitude of the auditory target used in the study.

#### Figure S2

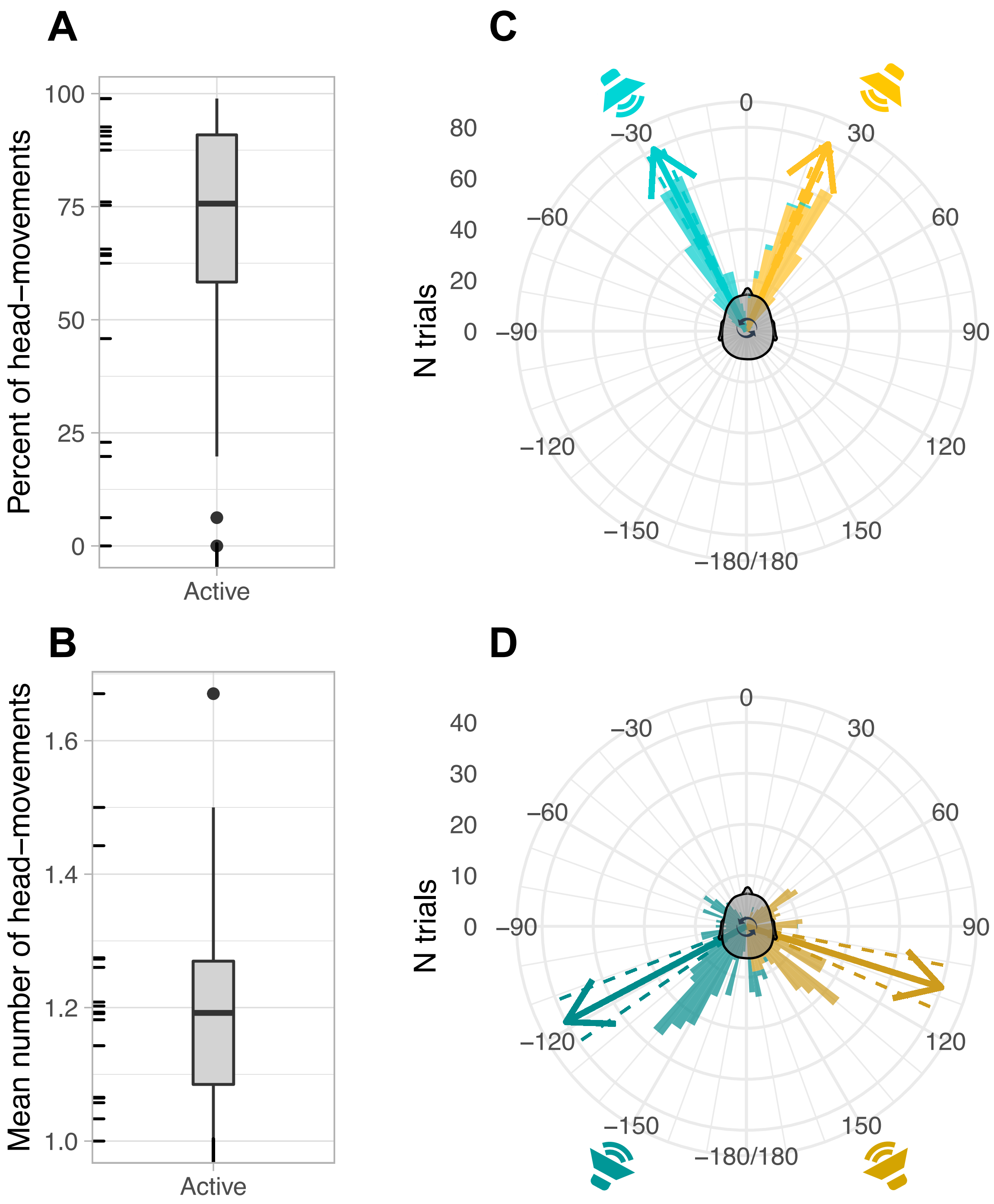


**Figure S2.** Head movements during sound in the active listening condition. (A) Boxplot of percentage head-movements. Note that two participants were identified as outliers (i.e., they fell outside 1.5 x interquartile range), made almost no head movement during the active listening condition and were thus excluded from subsequent analyses. (B) Boxplot of mean number of head movements once outliers are removed. (C-D) Polar histogram showing the distribution of head-movement responses for targets in front (C) and back (D) space. Arrow indicate mean head-movement direction, dashed lines indicate ± 1 SE.
